# NOTCH1 drives tumor plasticity and metastasis in hepatocellular carcinoma

**DOI:** 10.1101/2024.10.23.619856

**Authors:** Katherine E. Lindblad, Romain Donne, Ian Liebling, Erin Bresnahan, Marina Barcena-Varela, Anthony Lozano, Eric Park, Bruno Giotti, Olivia Burn, Maria I. Fiel, Clara Alsinet, Satdarshan P. Monga, Ruidong Xue, Jose Javier Bravo-Cordero, Alexander M. Tsankov, Amaia Lujambio

**Author notes:** **Corresponding author:** Amaia Lujambio, PhD. Department of Oncological Sciences, Liver Cancer Program, Division of Liver Diseases, Department of Medicine, Tisch Cancer Institute, Icahn School of Medicine at Mount Sinai, 1470 Madison Avenue Hess 6-111, New York, New York 10029 USA.

## Abstract

**Background & Aims:** Liver cancer, the third leading cause of cancer-related mortality worldwide, has two main subtypes: hepatocellular carcinoma (HCC), accounting the majority of the cases, and cholangiocarcinoma (CAA). Notch pathway primarily regulates the intrahepatic development of bile ducts, which are lined with cholangiocytes, but it can also be upregulated in 1/3 of HCCs. To better understand the role of NOTCH1 in HCC, we developed a novel mouse model driven by activated Notch1 intracellular domain (NICD1) and MYC overexpression in hepatocytes.

**Methods:** Using the hydrodynamic tail-vein injection method for establishing primary liver tumors, we generated a novel murine model of liver cancer harboring MYC overexpression and NOTCH1 activation. We characterized this model histopathologically as well as transcriptomically, utilizing both bulk and single cell RNA-sequencing. We also performed functional experiments using monoclonal antibodies.

**Results:** *MYC;NICD1* tumors displayed a combined HCC-CCA phenotype with temporal plasticity. At early time-points, histology was predominantly “cholangiocellular”, which then progressed to mainly “hepatocellular”. The “hepatocellular” component was enriched in mesenchymal genes and gave rise to lung metastasis. Metastatic cells were enriched in the TGFB and VEGF pathways and their inhibition significantly reduced the metastatic burden.

**Conclusions:** Our novel mouse model uncovered NOTCH1 as a driver of temporal plasticity and metastasis in HCC, the latter of which is, in part, mediated by angiogenesis and TGFß pathways.

**Impact and Implications:** This study develops a novel murine model of NOTCH1-driven liver cancer, an understudied oncogene in HCC. Using this model, we show that NOTCH1 drives plasticity in HCC and metastasis to the lungs that can be therapeutically targeted through inhibition of VEGF and TGFß pathways.

**Highlights:**

- NOTCH1 activation in combination with MYC overexpression drives combined HCC-CCA.
- NOTCH1 activation in hepatocytes drives temporal plasticity.
- NOTCH1 activation drives metastasis of HCC cells to the lungs, but not of CCA cells.
- Angiogenesis and TGFß pathways mediate NOTCH1-induced lung metastasis.

**Graphical abstract:** 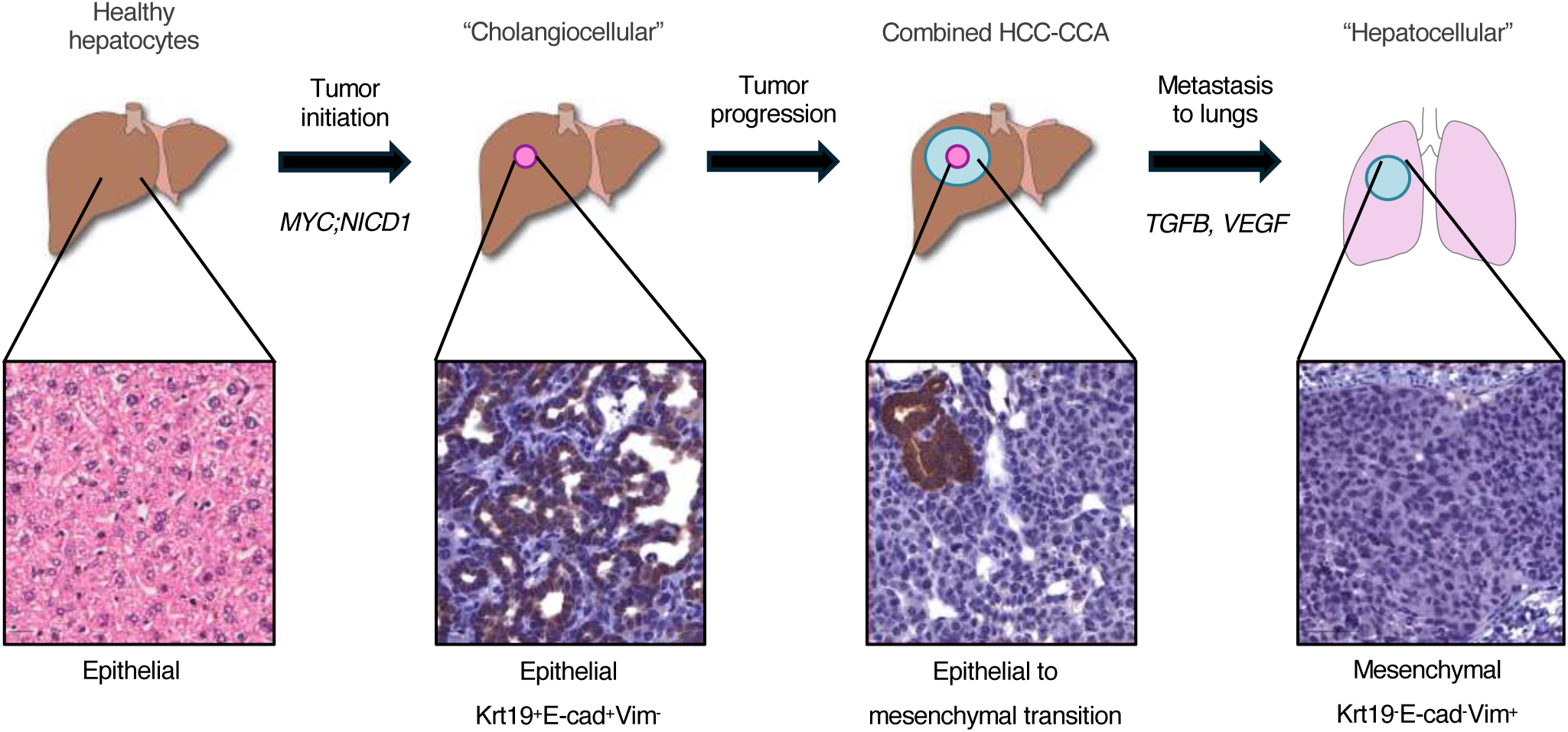

## Introduction

Notch signaling is a conserved inter-cellular communication pathway composed by four receptors (Notch1, 2, 3 and 4) and two ligand families [Jagged (JAG) 1 and 2 and Delta-like-ligand (Dll) 1, and 4] (1). The Notch signaling pathway involved ligand-induced activation of receptors, proteolytic cleavage, and subsequent translocation of the <u>N</u>otch Intra<u>C</u>ellular <u>D</u>omain (NICD) to the nucleus, where it functions as a transcriptional regulator (1). While Notch signaling may have a tumor suppressor role in certain contexts (2), Notch pathway activation has been described in around 30% of HCC patients (3, 4) and NOTCH1 has been validated as an oncogenic driver in mouse models of HCC (3, 4). We recently demonstrated that NOTCH1 activation plays a role in driving sexually dimorphic immune responses in HCC (5). However, its role in tumor progression and plasticity in HCC has not been explored.

Here, to elucidate the role of NOTCH1 activation in HCC tumor progression and plasticity, we characterized a novel mouse model of HCC driven by MYC expression and NOTCH1 activation (by overexpressing NICD1) that we recently generated (5) and demonstrated that NOTCH1 activation drives temporal tumor cell plasticity characterized by epithelial-to-mesenchymal transition (EMT) in vivo and metastasis of the mesenchymal component to the lungs. Interestingly, metastatic cells were enriched in the EMT, TGFB, and angiogenesis pathways, and targeting TGFB and VEGFR2 reduced the metastatic burden without affecting the growth of the primary tumor, shedding light into the mechanisms that dictate tumor progression and metastasis in HCC.

## Materials and methods

### Vector Use

All vector constructs were validated through Sanger sequencing and restriction enzyme digestion. The plasmids were kindly received from the following sources: *CMV-SB13,* Dr. Scott Lowe (MSKCC, New York); *pT3-EF1a-NICD1* (Addgene plasmid #46047) (6), Dr. Xin Chen (University of Hawaii Cancer Center). *pT3-EF1a-MYC-IRES-luciferase* (*MYC-luc*; Addgene plasmid #129775) and *pT3-EF1a-MYC-IRES-luciferase-OS* (*MYC-lucOS*; Addgene plasmid #129776) were generated as previously reported (7) as well as *px330-sg-p53* (8). We also generated the novel plasmid *pT3-EF1a-MYC-IRES-NICD1* by using the *pT3-EF1a-MYC-IRES-luc* and *pT3-EF1a-NICD1* plasmids.

### Hydrodynamic Tail Vein Injection

Vector production for use *in vivo* was performed either in house (QIAGEN plasmid *Plus*Mega kit; QIAGEN) or outsourced (Azenta). Nanodrop and agarose gel-based quantification methods were utilized to ensure equivalent DNA concentrations between batches to ensure reproducibility amongst experiments. Hydrodynamic tail vein injections were performed in mice 4-6 weeks of age, around 19 grams of bodyweight. Vectors were mixed at appropriate concentrations in 2mL of sterile 0.9% NaCl solution (Intermountain). Specifically, plasmid concentrations were prepared as follows: 11.4 μg of *pT3-EF1a-MYC-IRES-luciferase* (*MYC-luc*), 12 μg of *pT3-EF1a-MYC-IRES-luciferase-OS* (*MYC-lucOS*), 10 μg of *px330-sg-p53 (sg-p53),* 10 μg of *pT3-EF1a-NICD1* (*NICD*), and a 4:1 ratio of transposon to *SB13* transposase-encoding plasmid. A total volume of the saline/plasmid mix equivalent to 10% of body weight of each animal was injected into the lateral tail vein within 5-7 seconds (9). Two independent genetic alterations are required for malignant transformation in C57BL/6 mice (9), so only hepatocytes transfected with all three plasmids (transposon-based, CRISPR-based or transposon-based, and transposase-encoding) will have the potential to form tumors.

### Mice

Wild-type C57BL/6 mice were purchased from Envigo for each *in vivo* experiment unless otherwise specified. B6.Cg-Gt(ROSA)26Sortm9(CAG-tdTomato)Hze/J mice (Strain #007909) was obtained from Jackson. Each *in vivo* experiment was performed in accordance with the ISMMS Animal Care and Use Committee (protocol number IACUC-2014–0229). Each mouse utilized for *in vivo* experiments were maintained under specific pathogen-free conditions with *ad libitum* food and water. Prior to experimentation, all animals were assessed to ensure adequate health and acclimation to their environment. During necropsies of tumor-bearing mice, the liver, lungs, and spleen were collected and samples from each were frozen, embedded in OCT (Fisher), or formalin-fixed and paraffin-embedded.

### Bioluminescence imaging

Bioluminescence imaging was performed using an IVIS Spectrum system (Caliper LifeSciences; purchased with the support of NCRR S10-RR026561-01) to confirm equivalent transfection efficiency in a given experimental condition *in vivo*. 150 mg/kg D-luciferin (Thermo Scientific) was injected into mice intraperitoneally and resultant luciferase signal was assessed 5 minutes after administration of D-luciferin using Living Image software (Caliper LifeSciences). Prior to treatment initiation, mice were pre-stratified into cohorts with and equivalent average luciferase. Mice with an initial luciferase signal one log outside the average signal were excluded from the study.

### Immunohistochemistry

3 μm-thick sections from formalin-fixed and paraffin-embedded (FFPE) lung and liver tissues were utilized for all immunohistochemical staining. The protocol was performed as follows at ambient temperature unless otherwise indicated: Tissue deparaffinization in three 5-minute serial xylene treatments. Tissue re-hydration in decreasing ethanol concentrations (100%, 100%, 90%, and 80%) for 3 minutes each and then in distilled water for 5 minutes. Heated antigen retrieval in Target Retrieval Solution of the corresponding pH (Dako) and baked in a pressure cooker at 95°C for 15 minutes. Cooling of slides on ice followed by 3 washes in Tris-Buffered Saline pH 7.4 (TBS; Fisher). Blocking of endogenous peroxidase activity with 3% hydrogen peroxide (Sigma Aldrich) for 10 minutes, followed by three washes in TBS for 5 minutes. Protein blocking with Serum-Free Protein Block (Agilent) with appropriate serum for 30 minutes, followed by a brief rinse in TBS. Primary antibody staining in Background-Reducing Antibody Diluent (Agilent) with appropriate serum overnight at 4°C in a humid chamber. Washes, three times, 5-minute each, in TBS with 0.04% Tween20 (TBST) under agitation. Appropriate secondary antibody staining with appropriate serum for 30 minutes (ImmPRESS® HRP Universal Antibody Polymer Detection Kit; Vector Labs). Washes, three times, 5-minute each, in TBST under agitation. Antibody revelation using DAB Peroxidase Substrate Kit (Vector Labs) per manufacturer’s instructions. Counterstaining with Mayer’s Hemalum solution (Sigma-Aldrich). Tissue dehydration in serial ethanol (90%, 100%, 100%) for 15 seconds each and xylene twice for 5 minutes. Slide mounting with Permount Media (Fisher). Slide scanning at 20x resolution using Hamamatsu, NanoZoomer S60. The following antibodies were used (clone, company, dilution, antigen retrieval pH): MYC (Y69, abcam, 1:100, pH6), NOTCH1 (EP1238Y, abcam, 1:200, pH6), CK19 (EP1580Y, abcam, 1:200, pH6), Ki67 (D3B5, Cell Signaling, 1:200, pH6), VIMENTIN (D21H3, Cell Signaling, 1:200, pH6) and E-Cadherin (24E10, Cell Signaling, 1:200, pH6).

### RNA Extraction, RNA-Sequencing, and Analysis

Total RNA was isolated from 15-25 mg-sized, single, murine tumor nodules, which underwent mechanical digestion in cold Trizol® (Invitrogen), digestion with DNase I (Roche), and purification with RNeasy Kit (Qiagen). RNA sequencing (poly-A selected, PE150, 40 million reads) was outsourced through Genewiz. FeatureCounts (10) was used to measure read counts for each transcript and DESeq2 (11) was used to determine differentially expressed genes. A cut-off of 0.05 on adjusted p-value (corrected for multiple hypotheses testing) was used for creating gene lists. The GENI platform (https://www.shaullab.com/geni) (12) was used to perform GSEA. Student’s t test, corrected with Benjamini-Hochberg method, was applied for individually selected genes, to assess differences in gene expression values (TPM, transcripts per million). The files can be found at GEO through this link:

https://urldefense.proofpoint.com/v2/url?u=https3A__www.ncbi.nlm.nih.gov_geo_query_acc.cgi-3Facc3DGSE218377&#x0026;d=DwIBAg&#x0026;c=shNJtf5dKgNcPZ6Yh64b-ALLUrcfR4CCQkZVKC8w3o&#x0026;r=6ZW7xMgKmFhicC-cw4NTLDXXKHt5pu2yBJfk4_1T4c&#x0026;m=TqOE6bLGV2IIC5mdPSfuX_Nj-sBtHV40×5wKm_cXcULh8UE43TnzqA8tNFI4EZw&#x0026;s=qTvOGOb2Gx4rUJ3EMpb2axl5GXoV4NRv_n9_TBxz0rU&#x0026;e=

Enter token **cnsziucebbmfzqn** into the box.

### Tissue Dissociation for Single-Cell RNA Sequencing

At humane endpoint, mice were sacrificed, and the liver was perfused with at least 10 mL PBS+EDTA 2 mM through the inferior cava vein. Individual tumor nodules were extracted, weighed, and manually minced with sterile instruments. Visual inspection of selected tumor nodules was performed such to avoid healthy adjacent tissue as well as normalize for both size and level of expected necrosis across all samples. After manual dissociation of tumor nodules, enzymatic digestion was performed with 2.5mL enzymatic digestion mix (Miltenyi Biotech,Tumor dissociation Kit) in 1x DMEM in a gentleMACS C Tube (Miltenyi Biotech) and placed on gentleMACS Octo Dissociator (Miltenyi Biotech) with the “37°C_m_TDK1” program. Resultant tumor lysates were subjected to a 70-µm cell strainer (Miltenyi) and filtered cells were pelleted (300g for 5 min at 4°C). Red blood cell lysis was performed at 4°C using “Red blood cell lysis solution” (Miltenyi Biotech) for 2 min. Dead cells were removed using the “Dead Cell Removal” kit (Miltenyi Biotech) according to manufacturer protocols. CD45 positive cells were then separated from CD45 negative cells using the CD45 microbeads mouse kit (Miltenyi Biotech) at 4°C. Each fraction was manually counted by hemacytometer, then re-combined at 1:1 ratio, ensuring sufficient enrichment of immune cells to be sequenced from the bulk tumor samples. A cutoff for sample viability of >80% was used.

### Single-Cell RNA Sequencing Sample Processing

After cell isolation, single-cell RNA sequencing (scRNA-seq) was immediately performed by the HIMC core facility at the Icahn School of Medicine at Mount Sinai. To this end, single-cell suspensions were counted (Nexcelom Cellometer Auto 2000) and loaded on one lane of the 10x Genomics NextGem 5’v2.0 assay per the manufacturer’s protocol, with a targeted cell recovery of 10,000 cells per lane. Per the 10x Genomics protocol (https://assets.ctfassets.net/an68im79xiti/57JaTECQNBPSpyDz8oucdi/ced6aa8eaf73d6ee18dea8fdbd945faa/CG000331_Chromium_Next_GEM_Single_Cell_5-v2_UserGuide_RevD.pdf), gene expression (5’Gex) and TCR libraries were generated then quantified via Agilent Technologies 4200 Tapestation and KAPA library quantification kit (Roche). Targeted read depth included 25,000 reads per cell for gene expression libraries and 5,000 reads per cell for TCR libraries. Libraries were sequenced on Illumina NovaSeq S2 100 cycle kit with parameters set to 26×10×10×90 (R1xi7xi5xR2).

The files can be found at GEO through this link: https://urldefense.proofpoint.com/v2/url?u=https3A__www.ncbi.nlm.nih.gov_geo_query_acc.cgi-3Facc3DGSE218377&d=DwIBAg&c=shNJtf5dKgNcPZ6Yh64b-ALLUrcfR4CCQkZVKC8w3o&r=6ZW7xMgKmFhicC-cw4NTLDXXKHt5pu2yBJfk4_1T4c&m=TqOE6bLGV2IIC5mdPSfuX_Nj-sBtHV40×5wKm_cXcULh8UE43TnzqA8tNFI4EZw&s=qTvOGOb2Gx4rUJ3EMpb2axl5GXoV4NRv_n9_TBxz0rU&e=

### Single-Cell RNA Sequencing Data Pre-Processing, Dimensionality Reduction and Clustering

Libraries were processed with Cell Ranger v5.0.1, aligned to the mm10 reference genome, and then genes were filtered when expressing a cell number of <3. Seurat (v3.1.5) (13) was used to remove cells with UMIs sum (nFeatureRNA) of <200 and mitochondrion gene percentage of >10. 87,227 cells (+14,750 cells when lungs metastasis are included, n=2) passed the filter with an average of about 5800 cells/sample. Using Seurat, we performed dimensionality reduction and unsupervised clustering as follows: PCA was performed with all PCs loaded for UMAP dimensionality reduction. To identify clusters, these PCs were imported into FindClusters, an SNN graph-based clustering algorithm, with resolution of 2.5. Next, Wilcoxon Rank Sum test was performed to identify markers (FDR < 0.05) in each cluster (FindAllMarkers). After comparing with other datasets, cell type annotations were manually assigned based on these markers, which was further guided by functional enrichments using the Enrichr database (https://maayanlab.cloud/Enrichr/enrich, version June 8, 2023) (14). To identify subpopulations and remove doublets within these populations, each cluster was segregated and re-analyzed at higher resolution. For the single-cell TCR sequencing (scTCR-seq) analysis, T-cell receptor filtered contig files were generated from Cellranger v5.0.1 and then linked to individual cell barcode identifiers. TCR sequences were grouped into clonotypes, defined as groups of at least two cells sharing an identical TCR sequence, based on complementarity-determining region 3 (CDR3) sequence identity. Clonal diversity, expansion, and distribution within the sample was assessed. TCR data were integrated with other single-cell data (e.g., gene expression) to correlate TCR clonotypes with cellular phenotypes and functional states. RNA velocity analysis was conducted using the scVelo package (v0.3.2, Python v3.8.19). Aligned bam files from the Cell Ranger(v.8.0.1) outputs were used to generate HDF5-based loom files with the velocyto (v0.17.17) and loompy (v3.0.7) packages. The UMAP embedding of the single-cell data, along with metadata, was incorporated into the loom files to visualize the RNA velocity trajectory. Stochastic, deterministic, and dynamical modeling approaches were tested, with quality control plots of key marker genes guiding the selection of stochastic estimation as the final modeling method. The UMAP was generated using the scv.pl.velocity_embedding_stream() function from scVelo.

### Human HCC Sample Analysis

cBioPortal (15) TCGA LIHC dataset (n = 374) (16) or combined HCC-CCA and HCC cohort (n=311) (17) were used to obtain gene expression profiling data of HCC patients. Samples were stratified by *NOTCH1* expression (high = first tertile; low = second and third tertiles). Student’s t test, corrected with Benjamini-Hochberg, was performed to assess differences in gene expression values (TPM, transcripts per million). GSEA was performed using GENI platform (https://www.shaullab.com/geni) (12) or XENA (https://xena.ucsc.edu/) (18).

### Treatments

Day seven post-hydrodynamic delivery of the plasmids was chosen as the treatment initiation time point, one in which abundant microscopic lesions are present in the livers. Different treatment conditions were allocated to mice housed together to reduce cage-specific effects. All therapeutic monoclonal antibodies (mAbs) were administered intraperitoneally in 200 μL of the appropriate sterile antibody diluent (BioXcell). In the experiments with VEGFR2 and TGFß mAbs, three doses of either VEGFR2 (200 μg, clone DC101, BioXcell)/IgG1 (200 µg, clone HRPN, BioXcell) or TGFß (200 μg, clone 1S11.16.8, BioXcell)/IgG1 (200 µg, clone MOPC-21, BioXcell), were administered on days 7, 9, and 11.

### Statistical analysis

Mean ± standard deviation (SD) or median ± 95% confidence interval are used to convey data. Statistical significance was determined using the following: Mann-Whitney U test (when n<10 or non-normal distribution), Student’s t-test (n>10 and normal distribution), Wilcoxon test (paired comparisons), ANOVA or Kruskal-Wallis test (more than two groups), Kaplan-Meier and log-rank Mantle-Cox test (differences in survival), and chi-square test (association). For multiple comparisons, Benjamini-Hochberg (Student’s t-test) or Bonferroni (Mantle-Cox test) corrections were used. Significance values are as follows: *p<0.05, **p<0.01, ***p<0.001, ****<0.0001. Murine cohort size was determined based on the results of preliminary experiments. Allocation of mice into treatment cohorts was performed such that luciferase signal (transfection efficiency) was equivalent. Outcome assessment was not performed in a blinded manner. GraphPad Prism 9 software was used to generate graphs and statistical analysis.

## Results

### Characterization of a novel mouse model of HCC driven by NOTCH1 activation

To interrogate the role of NOTCH1 activation in HCC, which is overexpressed in one-third of HCC patients (3, 4), we recently generated a novel mouse model by performing hydrodynamic tail-vein injections (HDTVi) of transposon-based vectors expressing activated NOTCH1 (*pT3-EF1A-NICD1 (NICD)*, <u>N</u>otch Intra<u>C</u>ellular <u>D</u>omain, which contains the cytoplasmic domain initiating at amino acid 1753) (6), MYC (*pT3-EF1A-MYC*), and the SB13 transposase (*CMV-SB13*; enables integration and stable expression of both *MYC* and *NICD1*) into 6-week-old C57BL/6 mice (**Fig. S1A**) (5). We additionally performed HDTVi of a CRISPR-based vector expressing an sgRNA against p53 (*sg-p53*), *pT3-EF1A-MYC,* and *CMV-SB13* as a control for successful liver tumorigenesis (7) (**Fig. S1A**). Both *MYC;NICD1* and *MYC;sg-p53* mouse models displayed similar survival kinetics (35 and 39 days, respectively) (**Fig. 1A,B**) and tumor burden (quantified as liver weight) (**Fig. 1C**). However, NOTCH1 activation alone was not sufficient to cause death (**Fig. 1A**) or generate tumors in our model (**Fig. 1B; Fig. S1B**) within the study time-frame (120 days). However, NOTCH1 activation alone led to the formation of cystic nodules (**Fig. 1B; Fig. S1B**), in line with the role of NOTCH1 in driving biliary differentiation during development (19). Moreover, inhibition of NOTCH1 activation in *MYC;NICD1* tumors by concomitant overexpression of a dominant-negative form of NOTCH1 transcriptional activator RBPJ (dnRBPJ) led to a significant enhancement of overall survival in mice injected with *MYC;NICD1* (**Fig. 1D**), demonstrating the requirement of *MYC;NICD1* tumors on NOTCH1 activation.

**Fig. 1.**
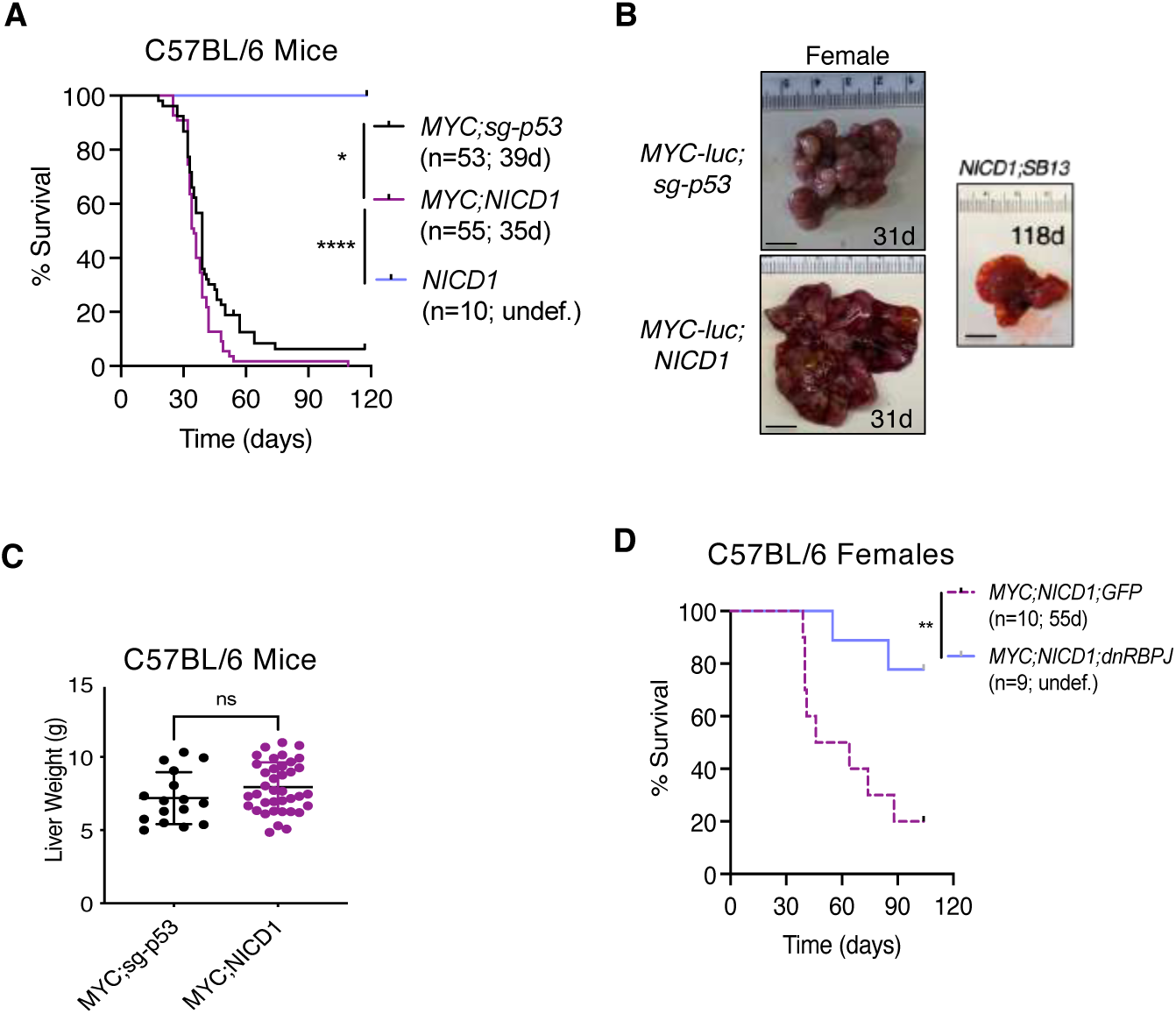
NOTCH1 activation in hepatocytes drives liver tumorigenesis. A,D) Survival curves in male and female C57BL/6 WT (wild-type) injected with *MYC;NICD1* (*MYC-lucOS;NICD1* or *MYC-luc;NICD1*) or *MYC;sg-p53 (MYC-lucOS;sg-p53* or *MYC-luc;sg-p53*) tumors or *NICD1;SB13* (A) or *MYC;NICD1;GFP* or *MYC;NICD1;dnRBPJ* (dnRBPJ; dominant negative form of RBPJ) tumors (D). Number of mice per group is shown as well as median survival in days (d). Log-rank Mantel-Cox test corrected with Bonferroni for multiple comparisons. B) Pictures of representative livers from A. The number indicates the number of days (d) from injection with the vectors to death for that particular mouse. Scale bar represents 1 cm. C) Liver weight in grams (g) at humane end point in mice harboring *MYC;sg-p53* or *MYC;NICD1* liver tumors. Student’s t test. Each dot represents one mouse.

### NOTCH1 activation drives combined HCC-CCA in mice

To dissect unique features of NOTCH1-driven tumors, we performed RNA-sequencing (RNA-seq) of bulk murine liver samples including healthy livers (n=6) and *MYC;NICD1* tumors (n=12), and compared these samples to the previously characterized *MYC;sg-p53* (n=17) and *MYC;CTNNB1* (n=12) endpoint tumors (7). Principal component analysis (PCA) demonstrated high similarity within each model (**Fig. S2A**): *MYC;CTNNB1* and *MYC;sg-p53* tumors showed a phenotype that was intermediate between healthy livers and *MYC;NICD1* tumors (PC1: 63.8%), while PC2 (15.9%) separated healthy tissue versus liver tumors. Unsupervised clustering based on the 2,500 most differentially expressed genes (DEGs) among samples clearly separated the four experimental groups (**Fig. 2A**). Among these 2,500 DEGs, NOTCH1 tumors (*MYC;NICD1)* were significantly enriched in Notch pathway genes (*Notch1*, *Notch3*, *Hes5*, *Hey1*, *Hey2*, *Jag1*, and *Spp1*) (4) compared to non-NOTCH1 tumors (*MYC;CTNNB1* and *MYC;sg-p53)* (**Fig. 2A,B**), highlighting successful pathway activation. Interestingly, genes associated with normal hepatocyte function (*Apob*, *Alb*, *Adh1*, *Hnf1a, Hnf4a*) were significantly decreased in NOTCH1-driven tumors while biliary-specific genes (*Krt19*, *Prdm5*, *Krt7*, *Epcam*) were significantly enriched (**Fig. 2A,B**), revealing a transcriptionally cholangiocarcinoma (CCA)-like phenotype (3, 20, 21). Single-sample Gene Set Enrichment Analysis (ssGSEA) (22) of Hallmarks datasets (23) and Hoshida HCC subclasses (24) revealed NOTCH1 tumors to be significantly enriched in “Notch_signaling” but significantly depleted in “adipogenesis”, “fatty_acid_metabolism”, “xenobiotic_metabolism”, and “bile_acid_metabolism” paythways (**Fig. S2B**), all of which are related to normal liver function. This confirms that NOTCH1-driven tumors are less hepatocytic than *p53*-deficient and *CTNNB1*-mutated murine tumors and is consistent with the role of NOTCH1 signaling in biliary differentiation (19). Accordingly, NOTCH1-driven tumors belonged to Hoshida S1 subclass (**Fig. S2B**), which is associated with moderately to poorly differentiated HCC tumors, poor survival, and NOTCH1 activation (4, 24). Interestingly, we have previously shown that *MYC;NICD1* tumors present sexual dimorphism in their immune responses (5); however, there were no differences in the transcript levels of genes related to Notch pathway, hepatocyte function, or biliary cells (**Fig. S2C**).

**Fig. 2.**
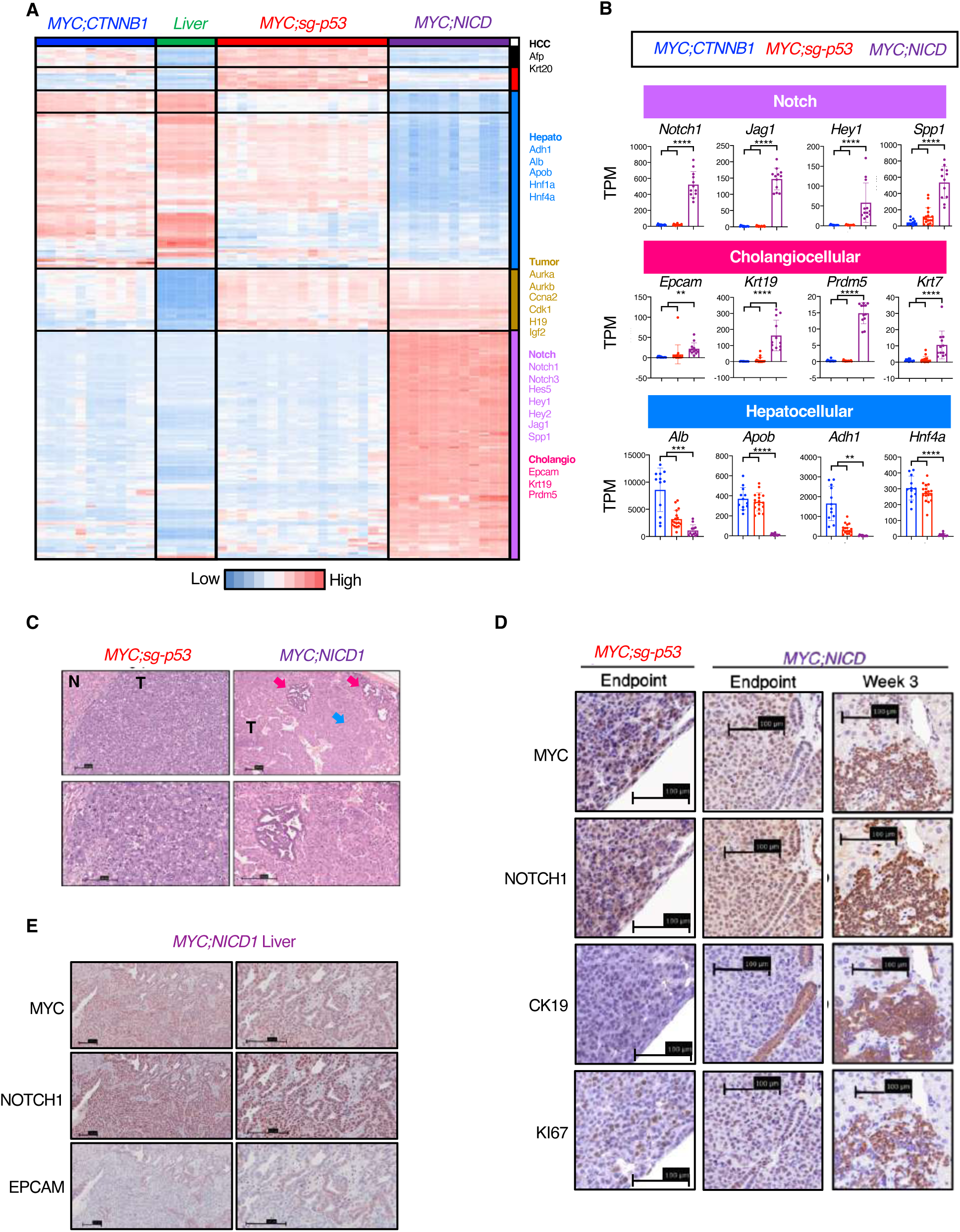
NOTCH1 activation in hepatocytes leads to combined HCC-CCA in mice. A) Heatmap of the top 2,500 differentially expressed genes in normal liver (n = 6) and *MYC;CTNNB1* (n = 12), *MYC;sg-p53* (n = 17), and *MYC;NICD1* (n = 20) tumors. Unsupervised clustering. Representative genes differentially expressed are highlighted on the right. B) Expression of representative genes in murine tumors. Student t test (Benjamini-Hochberg corrected) was performed comparing expression between non-NOTCH1 (*MYC;CTNNB1* and *MYC;sg-p53*) and NOTCH1 (*MYC;NICD1*) tumors. TPM, transcripts per million. Mean and standard deviation are shown. Each dot represents one sample. C) Hematoxylin-eosin staining of murine liver tumors sacrificed at humane endpoint. The bottom pictures are a magnification of the upper pictures. Blue arrow indicates “hepatocellular” region and pink arrows indicate “cholangiocellular” regions. T, tumor; N, normal liver. Scale bar represents 100 μm. D,E) Immunohistochemistry of MYC, NOTCH1, CK19, and KI67 (D), MYC, NOTCH1, and EPCAM (E) in representative murine tumors injected with the indicated vectors and sacrificed at humane endpoint or 3 weeks after HDTVI. Bar represents 100 μm. Ns, not significant. * p < 0.05. ** p < 0.01. *** p < 0.001. **** p < 0.0001.

Blinded histopathological assessment of the murine tumors by a liver pathologist revealed that *MYC;sg-p53* tumors were primarily of the macrotrabecular massive subtype and high-grade (**Fig. 2C**). In contrast, *MYC;NICD1* tumors showed a histology compatible with combined HCC-CCA (**Fig. 2C**); however, the “hepatocellular” regions were more abundant than CCA regions in tumors isolated at humane endpoint (**Fig. 2C**). Immunohistochemistry of CCA marker CK19 (cytokeratin 19, encoded by *Krt19* gene) showed staining in the CCA areas of *MYC;NICD1* tumors (**Fig. 2D**). Indeed, consecutive-slide analysis demonstrated that tumor cells could be either MYC^+^NOTCH1^+^CK19^+^ or MYC^+^NOTCH1^+^CK19^−^ (**Fig. 2D**). A similar expression pattern was presented by EPCAM, a typical marker of the CCA portion of combined HCC-CCAs (25) (**Fig. 2E**). In addition, all murine tumors were highly proliferative based on KI67 staining (**Fig. 2D**).

One of the drawbacks of the HDTVi of genetic elements is that while most transfected cells uptake all the plasmids in a given solution, there is also a possibility that a given hepatocyte may only uptake one or two plasmids. To confirm the combination of HCC-CCA was not due to the separate uptake of *MYC* or *NICD1* plasmids but their simultaneous expression in tumor cells, we generated a single plasmid encoding both genes (*pT3-EF1a-MYC-IRES-NICD1*) and performed HDTVi in combination with *CMV-SB13*. Histopathology on the resultant tumors revealed similar morphology to that of the HDTVi of separate plasmids, containing both HCC and CCA areas (**Fig. S2C**). Furthermore, injection of *pT3-EF1a-NICD1* and *CMV-SB13* plasmids alone led to the formation of cysts and a few non-malignant “cholangiocellular” areas (**Fig. S1B**), whereas *MYC;sg-p53* tumors maintained only HCC morphology (**Fig. 2C**) solidifying that NOTCH1 activation together with MYC overexpression is needed to promote tumor plasticity and combined HCC-CCA tumors in mice.

### NOTCH1 activation is associated with combined HCC-CCA in patients

To better understand the role of NOTCH1 in human HCC, we analyzed two patient cohorts, including the TCGA LIHC (Liver Hepatocellular) patient cohort (16) and one cohort of HCC and combined HCC-CCA patients (17). In both cohorts, there was a significant association between high *NOTCH1* expression (first tertile) and combined HCC-CCA tumor histology (**Fig. 3A,B**). Similar to the results observed in mice, in both patient cohorts, *NOTCH1*, Notch-targets such as *JAG1*, *HEY1, and SPP1*, as well as “Notch_signaling” signature were significantly upregulated in high-*NOTCH1* (first tertile) versus low-*NOTCH1* (second and third tertiles) tumors (**Fig. 3A,B; Fig. S3A,B**), validating the use of *NOTCH1* overexpression as a surrogate for NOTCH1 pathway activation in patients. In addition, CCA markers such as *EPCAM*, *KRT19*, *PRDM5*, and *KRT7* were significantly upregulated in high-*NOTCH1* tumors (**Fig. 3C,D**), further supporting the role of NOTCH1 in promoting a cholangio-like transcriptional phenotype (4). Interestingly, hepatocyte markers such as *ALB*, *APOB, ADH1,* and *HNF4A* were significantly downregulated in high-*NOTCH1* tumors in the combined HCC-CCA/HCC cohort (**Fig. 3D**) but not in the TCGA LIHC cohort, possibly because this cohort is enriched for HCCs (**Fig. 3A**). Nevertheless, in agreement with the murine data (**Fig. S2B**), high-*NOTCH1* tumors in both cohorts were significantly depleted in “adipogenesis”, “fatty_acid_metabolism”, “xenobiotic_metabolism”, and “bile_acid_metabolism” signatures (**Fig. S3A,B**), denoting that NOTCH1 drives loss of hepatic differentiation. Again mirroring the murine data, high-*NOTCH1* patient tumors were significantly enriched in the Hoshida Subclass 1 (S1) and conversely significantly depleted in the Hoshida Subclass 3 (S3), a subtype of well-differentiated tumors defined by activating *CTNNB1* mutations (**Fig. S3A,B**). Taken together, expression of *NOTCH1* in murine liver tumors leads to Notch pathway activation, upregulation of CCA markers, downregulation of HCC markers, and shows a combined HCC-CCA histopathological phenotype.

**Fig. 3.**
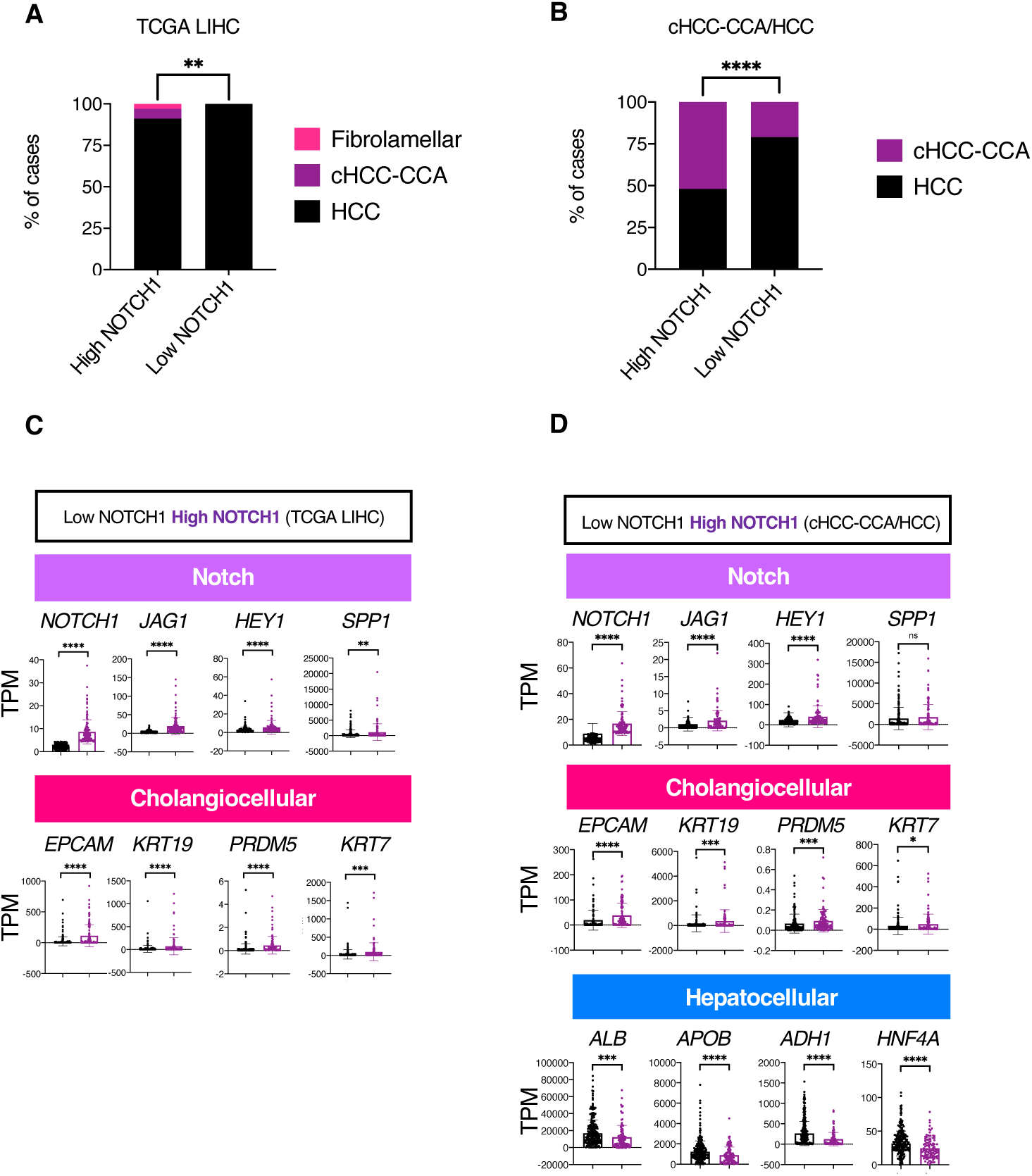
NOTCH1 activation is associated with combined HCC-CCA in patients. A,B) Association between high *NOTCH1* expression (first tertile) and HCC subtypes in the TCGA LIHC cohort (A) and cHCC-CCA/HCC cohort (B). Chi-squared test. C,D) Expression of representative genes in patient tumors. Student t test (Benjamini-Hochberg corrected) was performed comparing expression between low *NOTCH1* expression (second and third tertile) and high *NOTCH1* expression (first tertile) tumors. TPM, transcripts per million. Mean and standard deviation are shown. Each dot represents one sample.

### NOTCH1 activation promotes temporal plasticity

Of note, we found that at early time points, *MYC;NICD1* tumors primarily consisted of CCA areas whereas tumors isolated at humane endpoint primarily consisted of HCC areas (**Fig. 2D**), suggesting temporal plasticity. To further validate these findings, we performed single-cell RNA-sequencing (scRNAseq) analysis on *MYC;NICD1* tumors isolated at either an early time point (n = 2 females) or humane endpoint (n = 6 females) as well as endpoint *MYC;sg-p53* tumors (n=7), which could be clustered into 23 populations by UMAP (13) (**Fig. S4A,B**). Malignant cells, on the other hand, could be clearly separated into *MYC;sg-p53* and *MYC;NICD1* conditions (**Fig. S4C**). Further subclustering of *MYC;NICD1* malignant cells led to 16 groups of cells that were named according to the prevailing gene transcript or feature (**Fig. 4A,B**). The distribution of the distinct clusters was markedly different between early versus endpoint *MYC;NICD1* malignant cells (**Fig. 4A-C**). Early *MYC;NICD1* malignant cells were enriched in the subclusters “Spp1_Sox9” (MYC^low^ Notch1^high^ Spp1^high^ Epcam^high^ Sox9^high^ Mki67^high^) and “Spp1” (MYC^low^ Notch1^high^ Spp1^high^ Epcam^low^ Sox9^low^ Mki67^low^) while endpoint *MYC;NICD1* malignant cells were enriched in the subclusters “Malignant” (MYC^high^ Notch1^high^ Spp1^low^ Epcam^low^ Sox9^low^ Mki67^high^ Tagln^high^) and “Tagln” (MYC^high^ Notch1^high^ Spp1^low^ Epcam^low^ Sox9^low^ Mki67^low^ Tagln^high^) (**Fig. 4C; Fig. S4D**). This suggests that the “cholangiocellular” portion observed in the immunohistochemisty (**Fig. 2C**) may correspond with the “Spp1_Sox9” subcluster identified by scRNAseq and enriched in the early *MYC;NICD1* tumor samples (**Fig. 4C**). RNAvelocity analysis (26) suggested that most subclusters could be derived from this “Spp1_Sox9” subcluster (**Fig. 4D,E**). Indeed, “Spp1_Sox9” subcluster was enriched for transcripts related to liver progenitor cells (*Spp1, Epcam*, *Sox9, Krt8*, *Apoe*) (27) as well as a liver bipotent progenitor signature (**Fig. 4F,G; Fig. S4D,E**), confirming that *MYC;NICD1* tumors present a combined HCC-CCA phenotype and high plasticity.

**Fig. 4.**
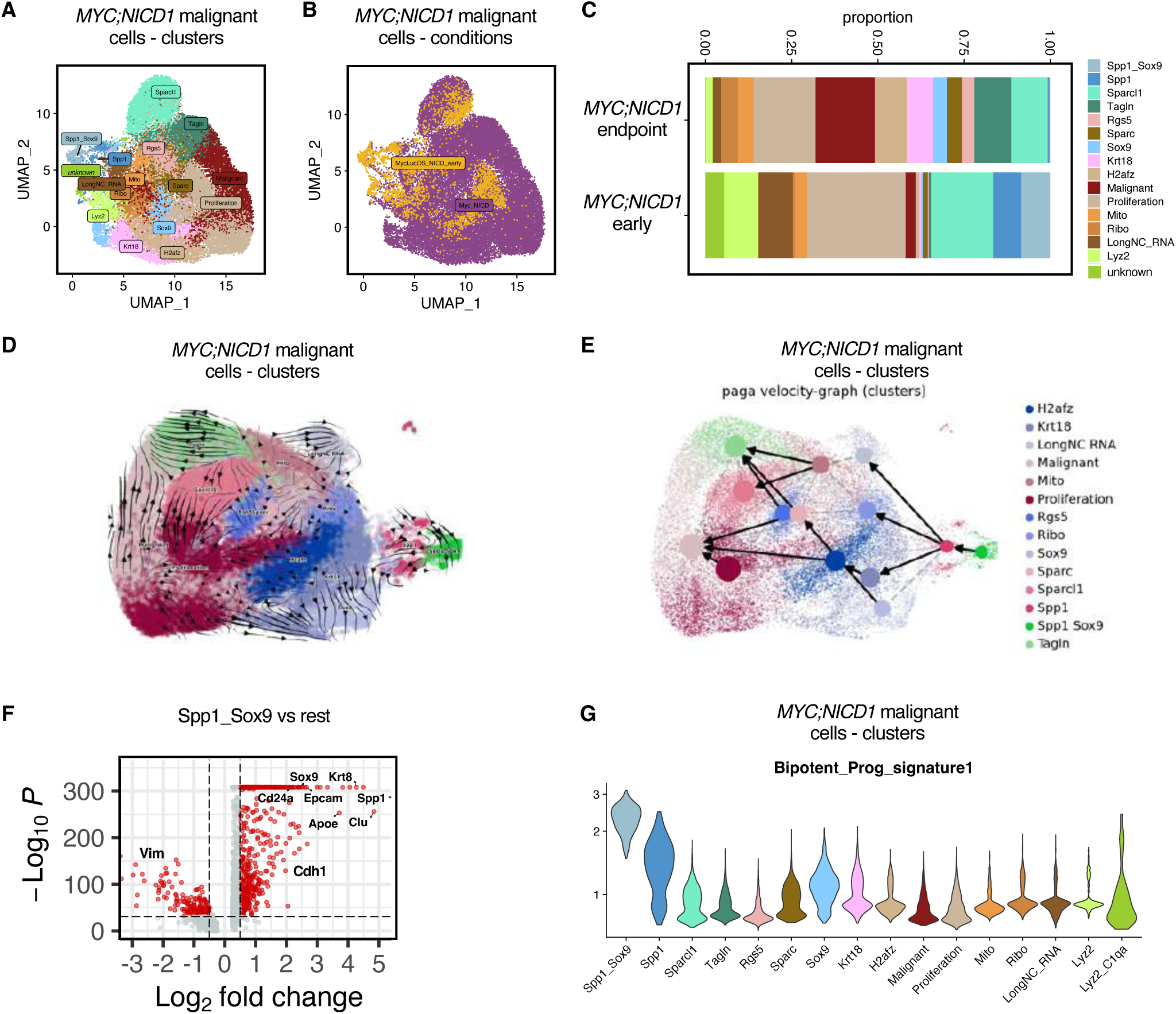
NOTCH1 activation promotes temporal plasticity. A,B) Single cell RNA sequencing (scRNAseq) UMAP plot depicting sub-clustered malignant cell population colored by tumor type (B) or sub-clusters (A). C) scRNAseq stacked bar plot displaying relative frequency of different subclusters within each tumor type shown in A. D,E) scRNAseq RNA velocity of malignant cells. Each dot represents a cell (D) or supercell (E). Cells are colored according to corresponding malignant sub-clusters. Arrows represent velocity that estimates extrapolated future cell states. F) scRNAseq volcano plot displaying top differentially expressed genes between *MYC;NICD1* progenitor cells (“Spp1_Sox9”) and *MYC;NICD1* non-progenitor cells shown in D,E. G) scRNAseq violin plots of progenitor gene signatures in different malignant sub-clusters.

### NOTCH1 activation is associated with EMT, TGFB, and angiogenesis signatures

Further analysis of the scRNAseq results indicated that the “Spp1_Sox9” subcluster was enriched in epithelial markers, such as E-cadherin (encoded by *Cdh1*), while the rest of the malignant cells were enriched in mesenchymal markers, such as Vimentin (encoded by *Vim*), suggesting that *MYC;NICD1* may undergo “Epithelial-to-Mesenchymal-Transition (EMT)” (**Fig. 4F**). Further analysis of the RNA-seq results showed that NOTCH1-driven tumors in both mice and patients are significantly enriched in the “Epithelial-to-Mesenchymal-Transition (EMT)”, “TGFβ”, and “Angiogenesis” Hallmark signatures (23) (**Fig. S2B, Fig. S3A,B**) – each of which are strongly associated with metastasis (28). Accordingly, EMT-related genes, (e.g. *Vim*, *Zeb2*, *Sparc,* and *Loxl1),* TGFβ pathway genes (e.g. *Tgfb1*, *Tgfb2, Tgfb1l1*, and *Bmp5)*, and angiogenesis-related genes (e.g. *Vegfa*, *Flt1* (*Vegfr1*), *Pdgfa*, and *Angpt2*) were significantly upregulated in NOTCH1-driven tumors (*MYC;NICD1)* compared to non-NOTCH1 tumors (*MYC;CTNNB1* and *MYC;sg-p53)* (**Fig. 5A**). Again, there were no significant differences in the transcript levels of genes related to EMT, TGFB, or angiogenesis when comparing tumors from females versus males (**Fig. S5B**).

**Fig. 5.**
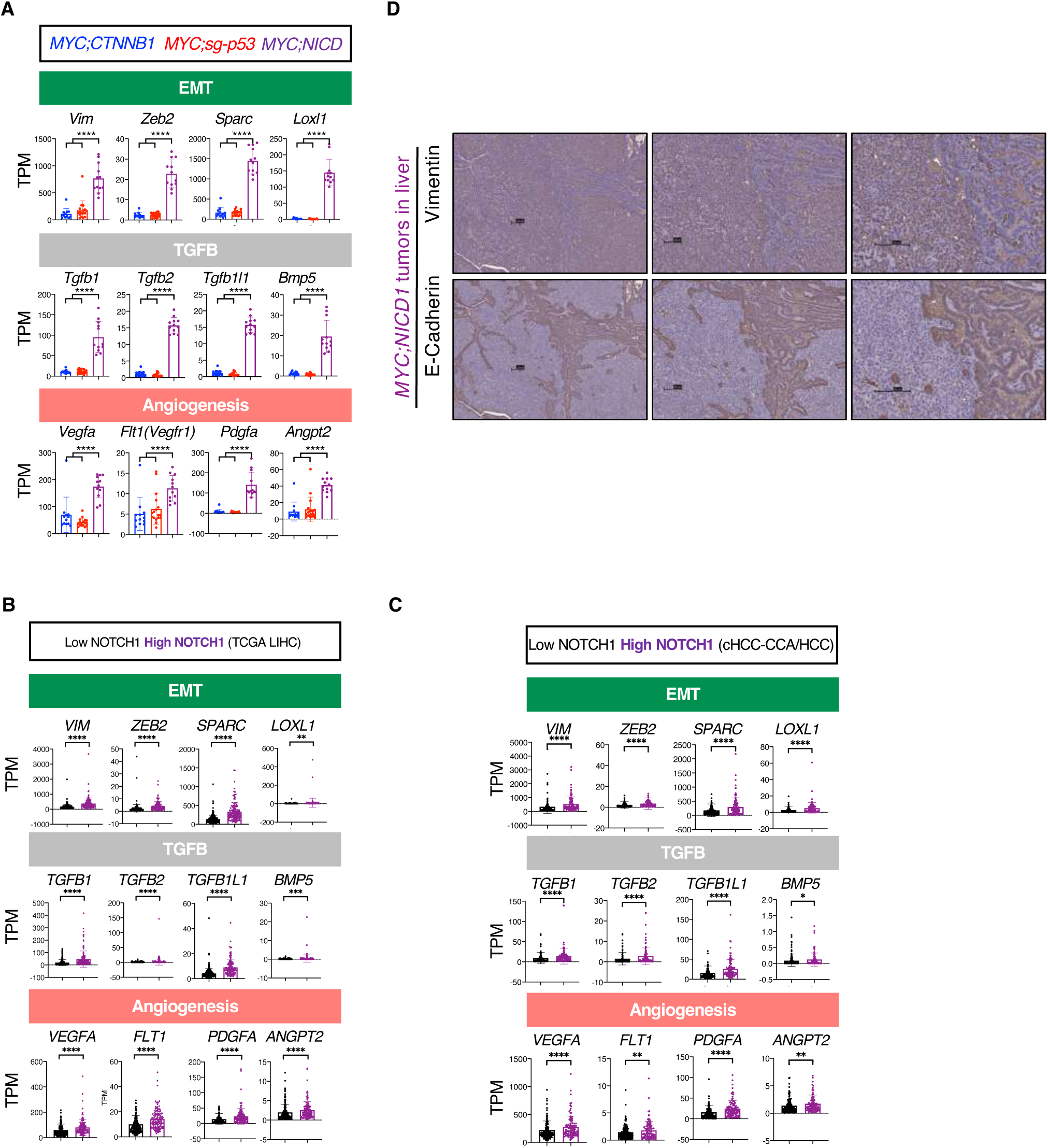
NOTCH1 activation is associated with upregulation of TGFB, EMT, and angiogenesis. A) Expression of representative genes in murine tumors. Student t test (Benjamini-Hochberg corrected) was performed comparing expression between non-NOTCH1 (*MYC;CTNNB1* and *MYC;sg-p53*) and NOTCH1 (*MYC;NICD1*) tumors. TPM, transcripts per million. Mean and standard deviation are shown. Each dot represents one sample. B,C) Expression of representative genes in patient tumors. Student t test (Benjamini-Hochberg corrected) was performed comparing expression between low *NOTCH1* expression (second and third tertile) and high *NOTCH1* expression (first tertile) tumors. TPM, transcripts per million. Mean and standard deviation are shown. Each dot represents one sample. D) Immunohistochemistry of Vimentin and E-Cadherin in a representative tumor from a *MYC;NICD1* mouse sacrificed at humane endpoint. The pictures to the right are a magnification of the pictures to the left. Bar represents 100 μm.

The association between NOTCH1 and EMT/TGFB/angiogenesis were confirmed in the two patient cohorts (16, 17), which showed that most of the same genes were significantly upregulated in high-*NOTCH1* tumors compared to low-*NOTCH1* tumors (**Fig. 5B,C**). Immunohistochemistry of E-cadherin (encoded by *Cdh1*; epithelial marker) and Vimentin (encoded by *Vim*; mesenchymal marker) further confirmed that NOTCH1-driven tumors presented pronounced EMT as tumor cells showed strong expression of Vimentin and low expression of E-cadherin (**Fig. 5D**). The pattern of E-cadherin and Vimentin expression was reversed in *MYC;sg-p53* tumors (**Fig. S5A**). Interestingly, the CCA portion of NOTCH1-driven tumors was negative for Vimentin and positive for E-cadherin (**Fig. 5D**), further confirming their progenitor state. Together, these results indicate that the “hepatocellular” component of NOTCH1-driven tumors is mesenchymal and could have metastatic potential.

### NOTCH1 activation promotes metastasis of “hepatocellular” cells to the lungs

Metastasis to the lungs occurs in approximately one-third of HCC patients (29) and, specifically, HCC tumors with Notch pathway activation have been associated with greater risk of metastasis to the lungs (30, 31). Detailed necropsy of the mice revealed metastasis to the lungs in the *MYC;NICD1* model in all the studied mice (30 out of 30) while only 1 out of 21 mice in the *MYC;sg-p53* model presented lung metastasis (**Fig. 6A,B**). The metastatic phenotype did not seem to be associated with the time between HDTVi and humane endpoint (median survival 39 days in the *MYC;sg-p53* model versus 35 days in the *MYC;NICD1* model) (**Fig. 1A**). Primary liver tumors are believed to be the principal cause of death in *MYC;NICD1* mice rather than metastatic tumor burden in the lungs, as we did not observe respiratory distress in most mice at endpoint, and *MYC;sg-p53* and *MYC;NICD1* mice developed liver tumors to a similar extent (quantified by liver weight as a surrogate for liver tumor burden; **Fig. 1C**). Hepatic origin of the primary and metastatic *MYC;NICD1* tumors was confirmed by lineage tracing and intravital two-photon microscopy (**Fig. 6C**). Here, *Rosa26-loxP-STOP-loxP-CAG-tdTomato* mice were injected with an adeno-associated virus expressing Cre recombinase under the liver-specific promoter Thyroxin Binding Globulin (*Tbg*) (*AAV8-Tbg-Cre*) to label hepatocytes with the fluorescent protein tdTomato (*Hep-tdTom*) (**Fig. 6C**). Four weeks later, *Hep-tdTom mice* were injected with *MYC-luc;NICD1* (**Fig. 6C**). At the time of death, *MYC-luc;NICD1* primary and metastatic tumors in *Hep-tdTom* mice were positive for tdTomato expression, confirming the hepatocytic origin (**Fig. 6C**). Interestingly, lung metastases were MYC^+^NOTCH1^+^CK19^−^ and presented a hepatocellular morphology, indicating that metastases are derived from MYC^+^NOTCH1^+^CK19^−^ primary tumor cells (**Fig. 6D,E**). Indeed, lung metastases were Vimentin positive and E-cadherin negative (**Fig. 6F)**, further demonstrating that only the HCC portion of the primary tumors leads to lung metastases.

**Fig. 6.**
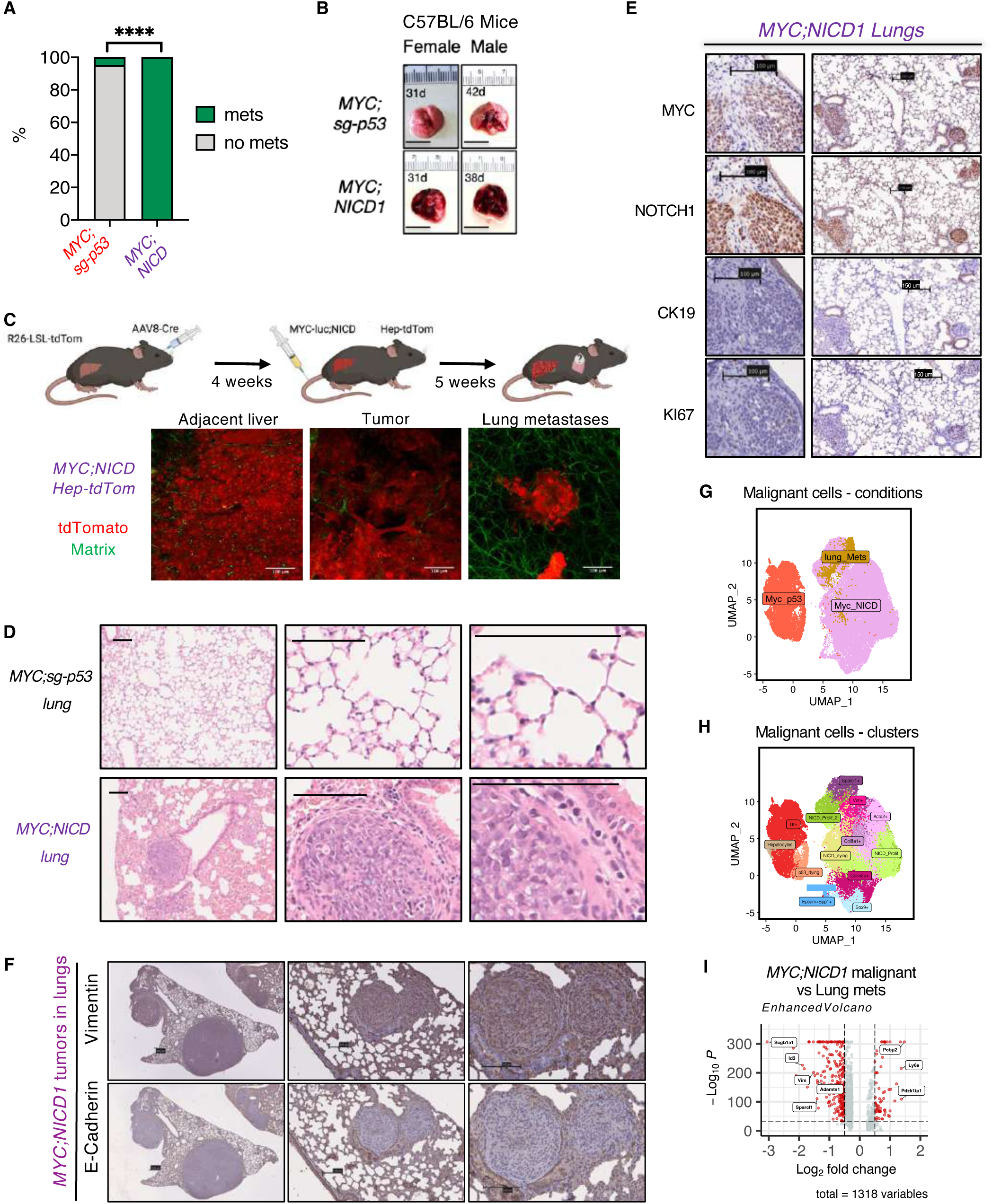
NOTCH1 activation drives metastasis to the lungs. A) Percentage of mice with lung metastases in the corresponding groups. Chi-square test. B) Images of representative lungs. The number indicates the number of days (d) from injection to death for that particular mouse. Scale bar represents 1 cm. C) Top, schematic of the experiment. Bottom, two-photon microscopy images of tdTomato (tdTom) liver tumors, adjacent liver tissue, and lung metastasis in a representative mouse. LSL, loxP-STOP-loxP; AAV8, adeno-associated virus 8. D) Hematoxylin-eosin staining of lungs isolated from tumor-bearing mice at humane endpoint. The pictures to the right are a magnification of the pictures to the left. Scale bar represents 100 μm. E,F) Immunohistochemistry of Vimentin and E-Cadherin (F) or MYC, NOTCH1, CK19, and KI67 (E) in representative samples from *MYC;NICD1* mice sacrificed at humane endpoint. The pictures to the right (F) or left (E) are a magnification of the pictures to the left or right, respectively. Bar represents 100 μm (E,F). G,H) Single cell RNA sequencing (scRNAseq) UMAP plot depicting sub-clustered *MYC;NICD1* malignant cell populations, including metastatic lung cells, colored by tumor type (G) or sub-clusters (H). I) scRNAseq volcano plot displaying top differentially expressed genes between *MYC;NICD1* liver tumor cells and *MYC;NICD1* metastatic lung cells. Ns, not significant. * p < 0.05. ** p < 0.01. *** p < 0.001. **** p < 0.0001.

To further understand the nature of the lung metastases, we performed scRNAseq of lung metastases from two female mice harboring *MYC;NICD1* tumors and added to the endpoint *MYC;sg-p53* and *MYC;NICD1* tumor samples shown in **Fig. 4**. The whole population clustered into 29 distinct populations by UMAP (**Fig. S6A-E**). Further sub-clustering of the malignant cells demonstrated that metastatic lung cells overlapped with *MYC;NICD1* tumor cells (**Fig. 6G-H; Fig. S6F-G**), implying that metastatic cells may already be present in the primary tumors as well as confirming their hepatic origin. Accordingly, a low number of genes were differentially expressed in metastatic lung cells when compared to the primary tumor cells from *MYC;NICD1* tumors (**Fig. 6I**), further indicating that some primary tumor cells already have metastatic potential. Among the genes that were upregulated in metastatic lung cells were *Adamts1* (metalloproteinase) (32) and *Sparcl1* (extracellular matrix binding), which could potentially be involved in adaptation to growth in the lungs (**Fig. 6I**). Indeed, metastatic lung cells were enriched in the Sparcl1 subcluster (**Fig. S6G**). Moreover, metastatic lung cells did not overlap with *MYC;NICD1* progenitor tumor cells (**Fig. 6G,H**), confirming that only the cells from the “hepatocellular” and not the “cholangiocellular” compartment metastasize to the lungs. Taken together, NOTCH1 activation promotes metastasis to the lungs in this novel mouse model of liver cancer.

### Therapeutic inhibition of TGFß and VEGF ameliorates metastatic lung burden

The EMT, angiogenesis, and TGFß pathways were further enriched in the metastatic lung cells compared to the primary *MYC;NICD1* tumor cells (**Fig. 7A-F**), underscoring the importance of these pathways in the mechanism of metastasis to the lungs. We next asked whether therapeutically targeting these pathways may lead to an increase in survival and/or reduction in metastatic tumor burden. Using monoclonal antibodies targeting either TGFß or VEGFR2 alone or in combination (**Fig. S7A**) had no effect on survival outcome (**Fig. 7G**) or tumor burden (**Fig. S7B**) in mice harboring *MYC;NICD1* tumors compared with their IgG counterparts. However, a significant reduction in metastatic tumor burden was observed in mice receiving anti-TGFß, anti-VEGFR2, or the combination thereof compared to IgG controls (**Fig. 7H,I**). Interestingly, the combination of both anti-TGFß and anti-VEGFR2 did not lead to a significant reduction in metastatic tumor burden compared with the single agent controls, suggesting a lack of synergy between the two therapeutics. These results further highlight the importance of TGFß and VEGF pathways in mediating NOTCH1-driven lung metastasis in liver cancer and further open the avenue for therapeutic interventions in patients with metastatic liver cancer.

**Fig. 7.**
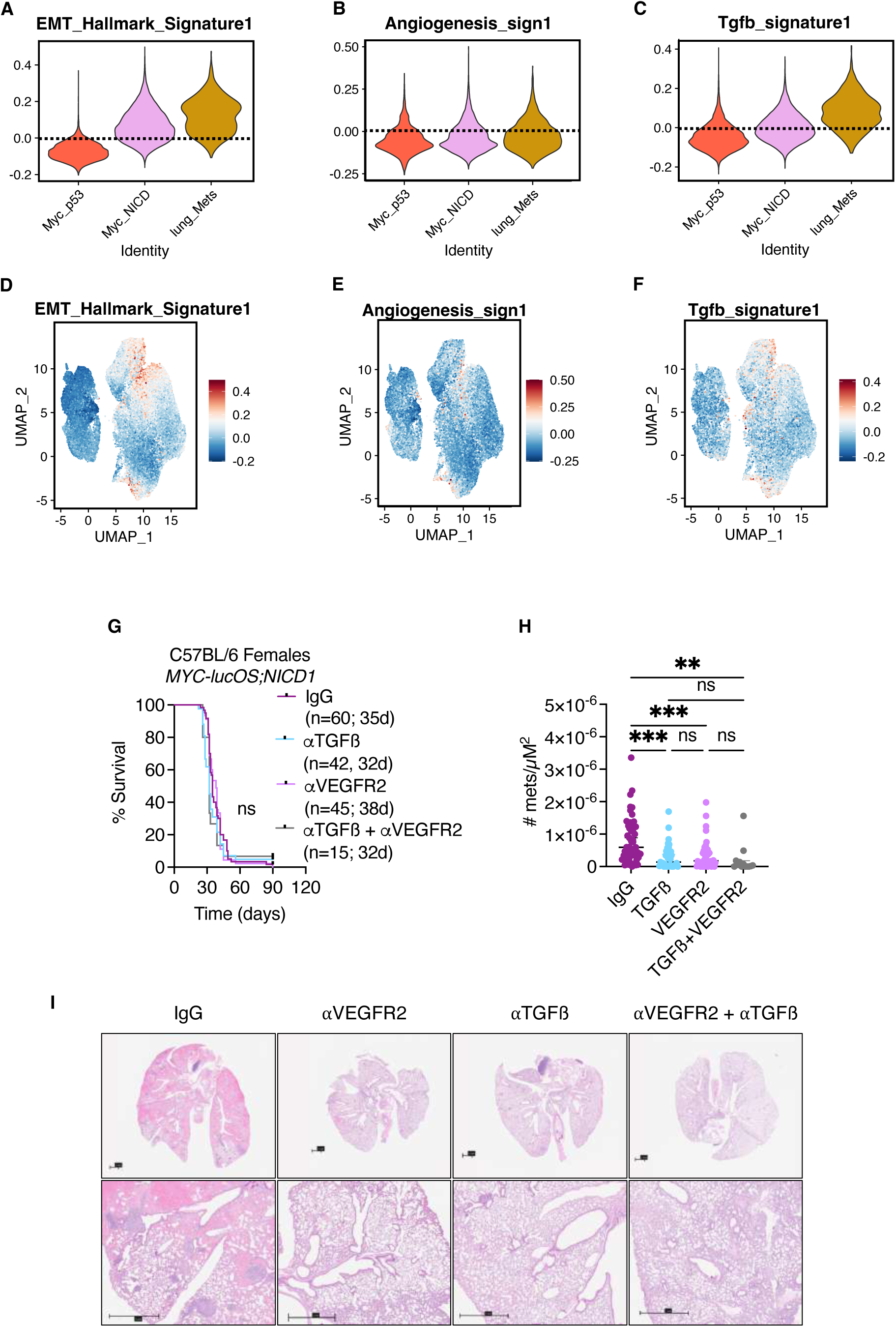
Lung metastases are inhibited by TGFß and VEGF blockade. A-C) scRNAseq violin plots displaying expression level of EMT (A), angiogenesis (B), and TGFß (C) signatures across each cluster shown in Fig. 6G. D-F) Single cell RNA sequencing (scRNAseq) UMAP plot depicting sub-clustered malignant cell populations shown in Fig. 3G displaying relative expression of EMT (D), angiogenesis (E), and TGFß (F) signatures within each population. G) Survival curve in C57BL/6 WT (wild-type) female mice. Number of mice per group is shown as well as median survival in days (d). Log-rank Mantel-Cox test corrected with Bonferroni for multiple comparisons. H) Number of metastases normalized per μm^2^ of tissue. One-way ANOVA with multiple comparisons to IgG control was performed. Each dot represents one mouse. I) Representative hematoxylin-eosin staining of lungs isolated at humane endpoint in mice treated with the indicated treatments. The pictures on the bottom are a magnification of the pictures on the top. Bar represents 1 mm. Ns, not significant. * p < 0.05. ** p < 0.01. *** p < 0.001. **** p < 0.0001.

## Discussion

While NOTCH1 can potentially play a role as a tumor suppressor in HCC (2), it predominantly acts as an oncogene in HCC (3, 33). In previous studies, NOTCH1 activation, through the expression of NICD1 in murine hepatocytes, was shown to promote HCC, either alone (3) or in combination with non-alcoholic steatohepatitis (4). In our study, we demonstrate that NOTCH1 activation cooperates with MYC overexpression to generate HCCs that harbor combined HCC-CCA histopathology and can be transcriptomically defined as cholangio-like tumors. The most interesting finding is the temporal plasticity that we observe in *MYC;NICD1* tumors where in early stages of tumorigenesis, the “cholangiocellular” compartment is more prominent, while at later stages, the “hepatocellular” portion takes over. Single-RNA sequencing studies of early and late tumor samples revealed that the “cholangiocellular” cells are of progenitor nature while the “hepatocellular” cells are mesenchymal. RNA velocity analysis suggests that these “cholangiocellular” cells would lead to the “hepatocellular” cells. Future studies will unveil the mechanism behind this temporal plasticity. To our knowledge, the *MYC;NICD1* mouse model is unique at displaying EMT in such a defined way.

In addition to tumor plasticity, cooperation between MYC and NOTCH1 also contributes to metastasis, an important step of HCC pathogenesis, implying that the *MYC;NICD1* model may represent a tractable platform for studying these two important cancer processes. Of note, only “hepatocellular” cells, which present a mesenchymal phenotype, but not “cholangiocellular” cells, which are epithelial, are able to metastasize. Cells that metastasize to the lungs seem to be already present in the primary tumors, at early and late time stages, and present higher mesenchymal features as well as enrichment of the TGFB and angiogenesis pathways compared to other NOTCH1-expressing tumor cells. Accordingly, inhibition of TGFB or angiogenesis reduced the metastatic burden without affecting the primary tumor burden, indicating that these two pathways have exclusive roles in metastasis in our model. Taken together, our murine studies demonstrate that NOTCH1 is a critical oncogene in HCC that contributes to tumor plasticity and metastasis, in addition to tumor initiation. We further validated these phenotypes in patient samples. Importantly, our studies highlight the value of rationally designed mouse models that can lead to a better understanding of tumor biology.

## Authors’ Contributions

**Conception and design:** K.E. Lindblad, A. Lujambio.

**Development of methodology:** K.E. Lindblad, A. Lujambio.

**Acquisition of data (provided animals, acquired and managed patients, provided facilities, etc.):** K.E. Lindblad, R. Donne, I. Liebling, M. Barcena-Varela, A. Lozano, E. Bresnahan, O. Burn. **Analysis and interpretation of data (e.g., statistical analysis, biostatistics, computational analysis):** K.E. Lindblad, R. Donne, I. Liebling, E. Bresnahan, M. Barcena-Varela, A. Lozano, E. Park, B. Giotti, O. Burn, M.I. Fiel, N. Param, C. Alsinet, S.P. Monga, R. Xue J.J. Bravo-Cordero, A.M. Tsankov, A. Lujambio

**Writing, review, and/or revision of the manuscript:** K.E. Lindblad, A. Lujambio with comments from the rest of the authors.

**Administrative, technical, or material support (i.e., reporting or organizing data, constructing databases):** K.E. Lindblad, A. Lujambio.

**Study supervision:** K.E. Lindblad, A. Lujambio.

## Acknowledgments

We thank Drs. Scott Lowe, Xin Chen, Tyler Jacks, and Feng Zhang for access to plasmids. We thank the Icahn School of Medicine at Mount Sinai (ISMMS) Center for Comparative Medicine and Surgery (CCMS), the ISMMS Translational and Molecular Imaging Institute (TMII) Imaging Core, the Tisch Cancer Institute Flow Cytometry Shared Resource Facility, the Tisch Cancer Institute Microscopy Shared Resource, the Oncological Sciences Histology Shared Resource Facility, the Oncological Sciences ImmunoStaining Shared Resource Facility, and the Microscopy and Advanced Bioimaging Core Facility at Mount Sinai. We thank Drs. Scott L. Friedman (Sinai), Jeremiah Faith (Sinai), Augusto Villanueva (Sinai), Brian Brown (Sinai), Cecilia Berin (Sinai), David Dominguez-Sola (Sinai), Ming Li (MSKCC), and Changyu Zhu (MSKCC) for insightful comments. Part of this manuscript was used in a Dissertation Thesis by Katherine E. Lindblad for the Graduate School of Biomedical Sciences at the Icahn School of Medicine at Mount Sinai.

## SUPPLEMENTARY LEGENDS

**Fig. S1.**
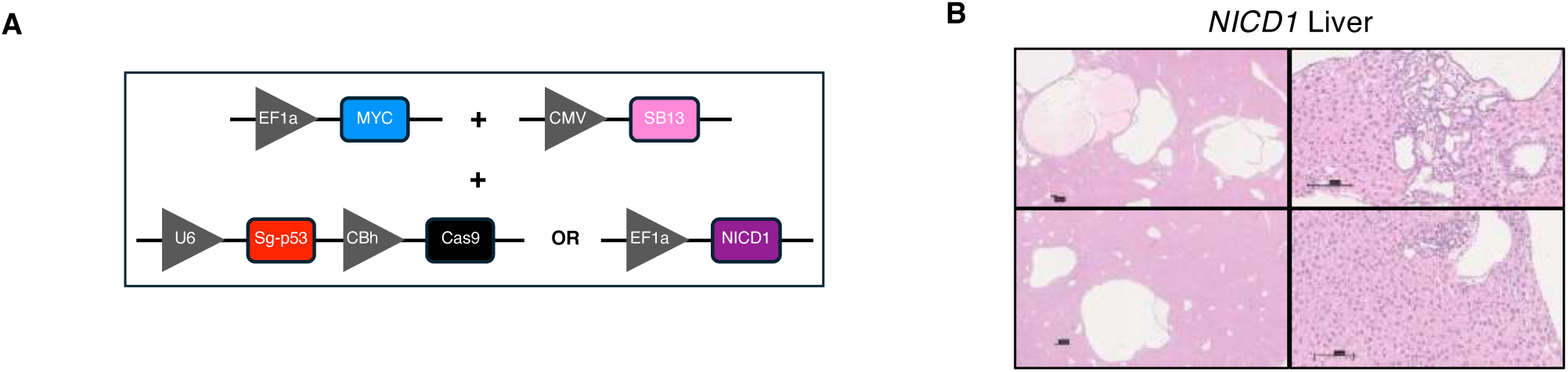
NOTCH1 activation in hepatocytes drives liver tumorigenesis. A) Schematic of vectors injected into mice. B) Hematoxylin-eosin staining of two representative murine livers. The picture on the bottom is a magnification of the picture on the upper part. Bar represents 100 μm.

**Fig. S2.**
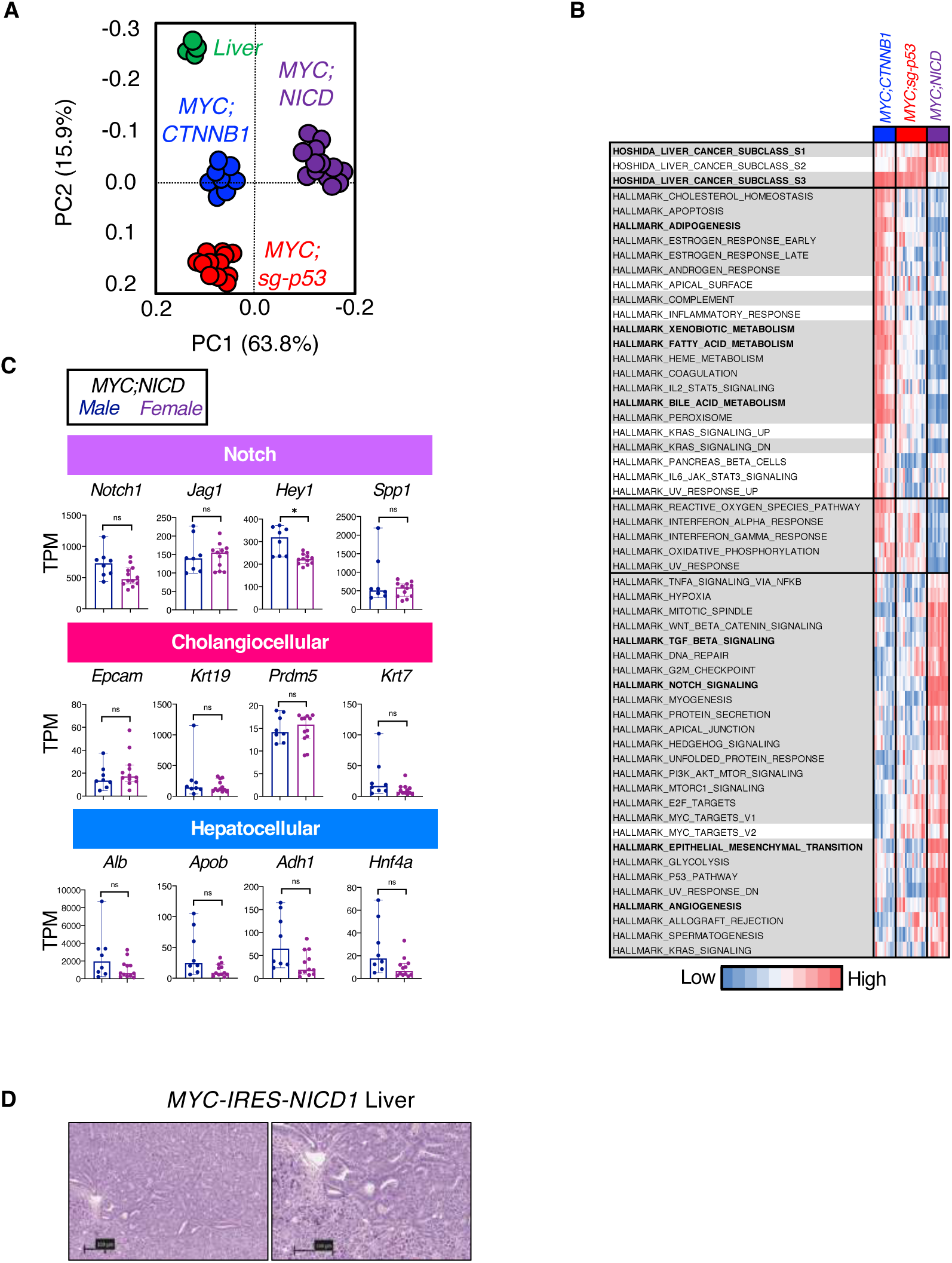
NOTCH1 activation in hepatocytes leads to combined HCC-CCA in mice. A) Principal Component Analysis (PCA) of normal liver (n = 6) and *MYC;CTNNB1* (n = 12), *MYC;sg-p53* (n = 17), and *MYC;NICD1* (n = 20) endpoint tumors. B) Heatmap representing the results from ssGSEA analysis [5] in murine tumors for the Hallmarks and Hoshida gene signatures. Significant gene signatures are highlighted in grey. Student t test (Benjamini-Hochberg corrected) between non-NOTCH1 (*MYC;CTNNB1* and *MYC;sg-p53*) and NOTCH1 (*MYC;NICD1*) murine tumors. Signatures mentioned in the text are highlighted in bold. C) Expression of representative genes in murine tumors. Student t test (Benjamini-Hochberg corrected) was performed comparing expression between female and male *MYC;NICD1* tumors. TPM, transcripts per million. Mean and standard deviation are shown. Each dot represents one sample. D) Hematoxylin-eosin staining of a representative murine tumor. The picture on the right is a magnification of the picture on the left. Bar represents 100 μm.

**Fig. S3.**
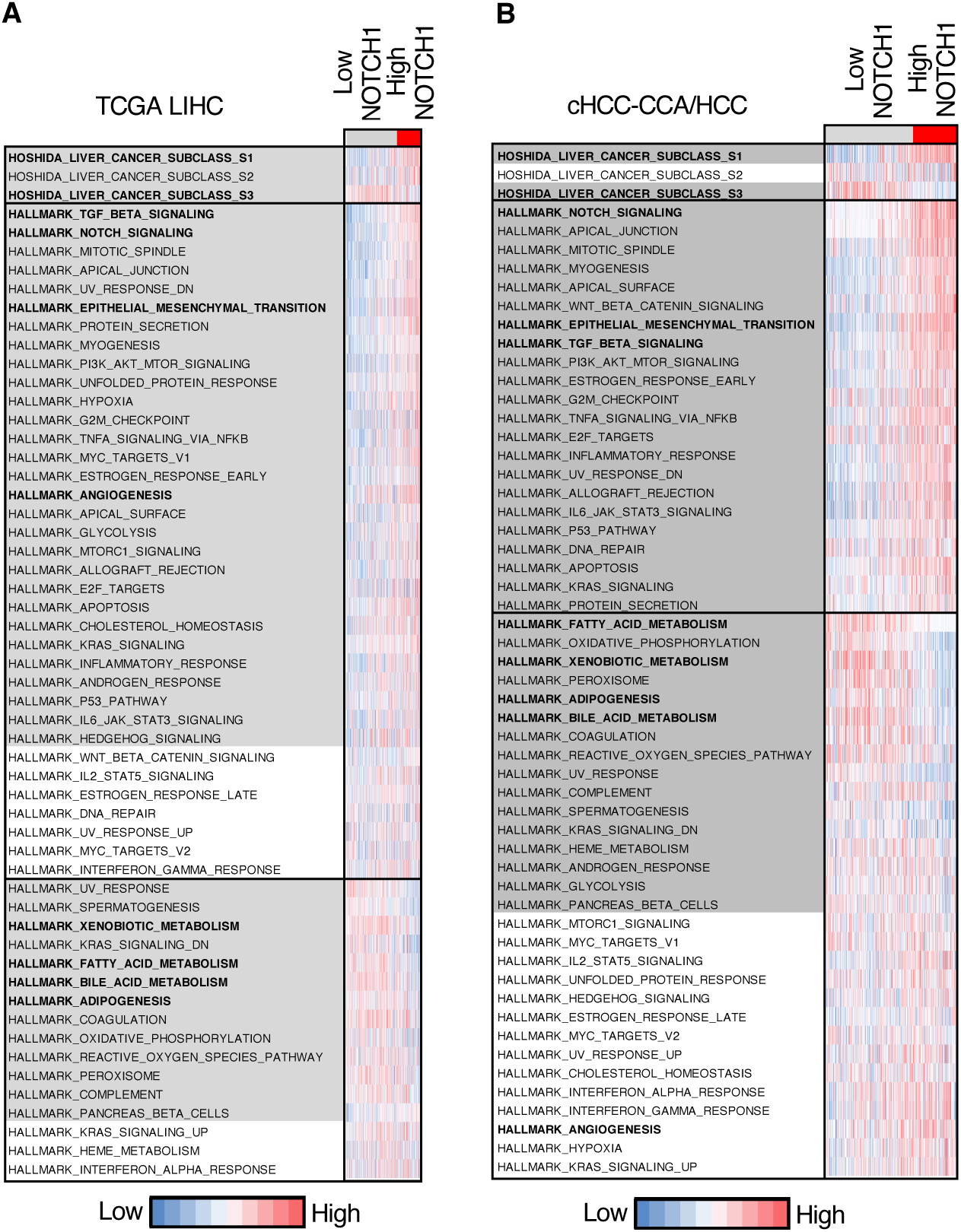
NOTCH1 activation is associated with combined HCC-CCA in patients. A,B) Heatmap representing the results from ssGSEA analysis [5] in TCGA LIHC cohort (A) or combined HCC-CCA (cHCC-CCA)/HCC cohort (B) for the Hallmarks and Hoshida gene signatures. Significant gene signatures are highlighted in grey. Student t test (Benjamini-Hochberg corrected) between low-*NOTCH1* (second and third tertiles) and high-*NOTCH1* (first tertile) tumors was performed. Signatures mentioned in the text are highlighted in bold.

**Fig. S4.**
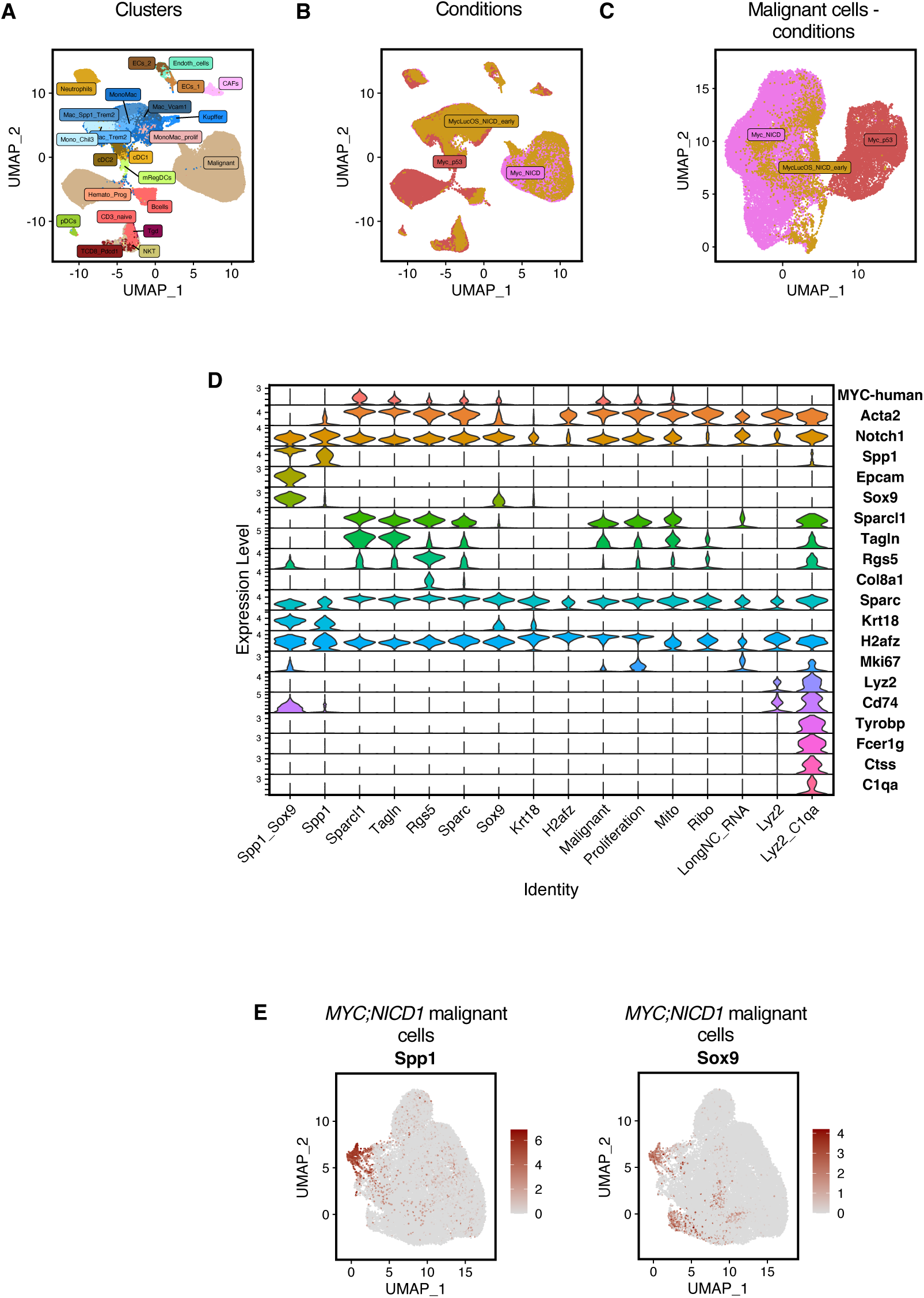
NOTCH1 activation promotes temporal plasticity. A-C) Single cell RNA sequencing (scRNAseq) UMAP plot of liver tumors analyzed (n = 7 female *MYC;sg-p53*, n = 6 female *MYC;NICD1*, n = 2 early time-point *MYC;NICD1*) by single cell RNA sequencing (scRNAseq) showing clusters by cell type into 23 unique populations (A) or tumor conditions (B). Sub-clustered malignant cells are shown by tumor type (C). D) scRNAseq violin plot displaying expression level of cell identity genes in each of the clusters of *MYC;NICD1* malignant cells. E) Single-cell RNA sequencing (scRNAseq) UMAP plot depicting sub-clustered *MYC;NICD1* malignant cell population colored by expression levels of indicated genes.

**Fig. S5.**
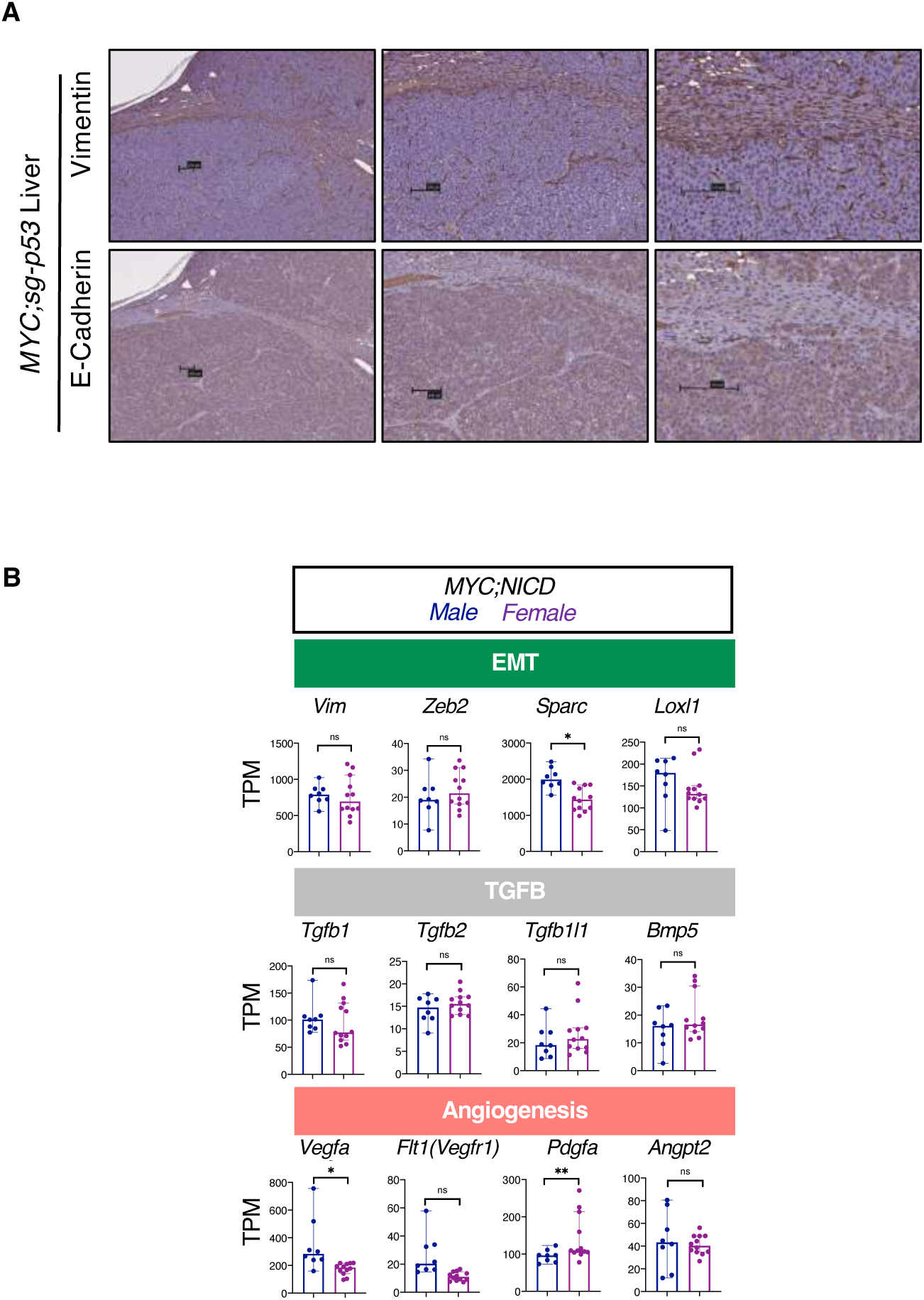
NOTCH1 activation is associated with upregulation of TGFB, EMT, and angiogenesis. A) Immunohistochemistry of Vimentin and E-Cadherin in a representative tumor from a *MYC;sg-p53* mouse sacrificed at humane endpoint. The pictures to the right are a magnification of the pictures to the left. Bar represents 100 μm. B) Expression of representative genes in murine tumors. Student t test (Benjamini-Hochberg corrected) was performed comparing expression between female and male *MYC;NICD1* tumors. TPM, transcripts per million. Mean and standard deviation are shown. Each dot represents one sample.

**Fig. S6.**
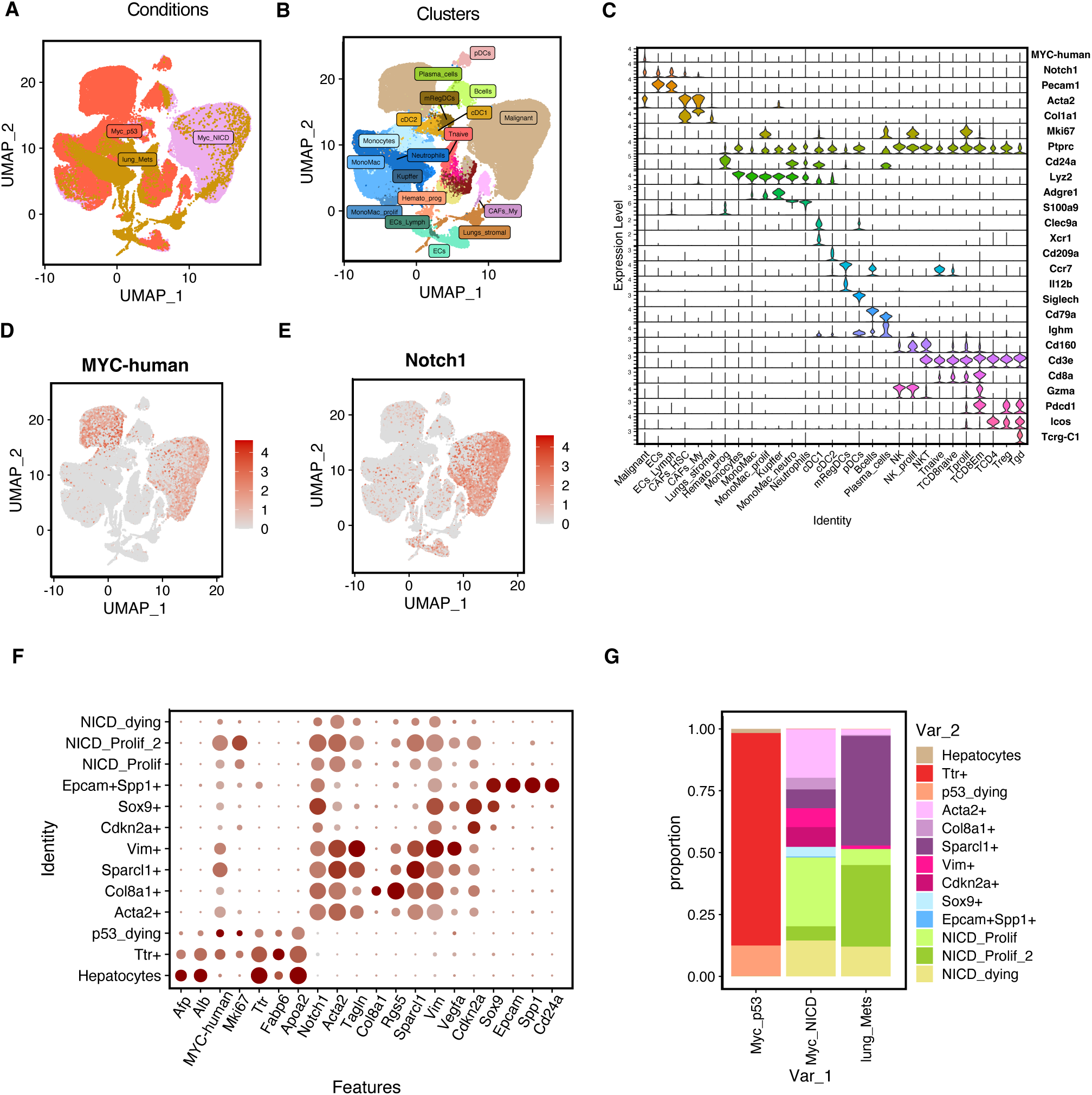
NOTCH1 activation drives metastasis to the lungs. A,B,D,E) Global UMAP plot of liver tumors analyzed (n = 7 female *MYC;sg-p53*, n = 6 female *MYC;NICD1*, n = 2 lung *MYC;NICD1* metastases) by single cell RNA sequencing (scRNAseq) colored by tumor type (A), clustered by cell type into unique populations (B), or displaying relative expression of indicated gene transcripts within each population (D,E). C) scRNAseq violin plot displaying expression level of cell identity genes in each of the cell clusters identified in B. F) scRNAseq dot plot displaying relative expression of key transcripts within the malignant sub-clusters. G) scRNAseq stacked bar plot displaying relative frequency of different sub-clusters within each condition.

**Fig. S7.**
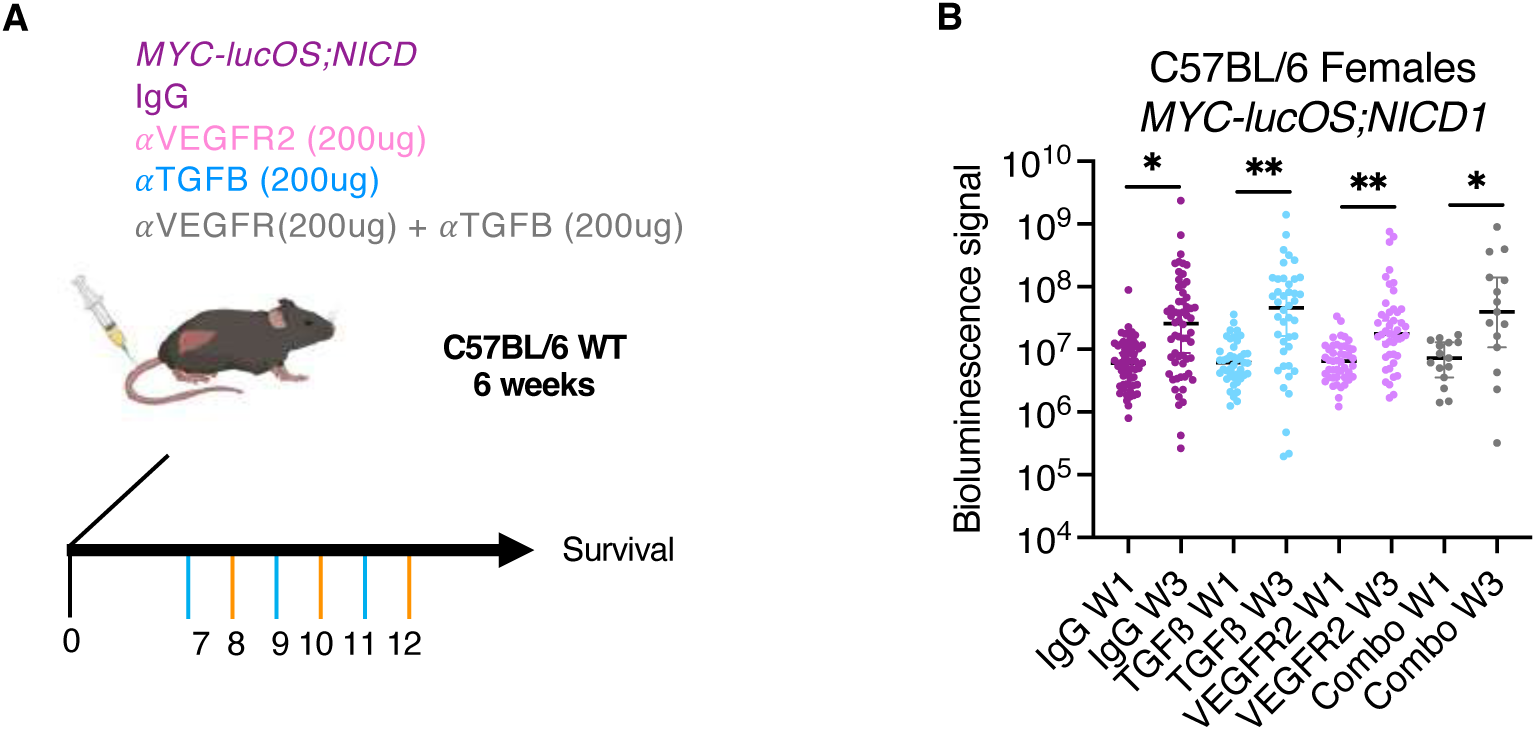
Lung metastases are inhibited by TGFß and VEGF blockade. A) Schematic of experiment shown in Fig. 7G. B) Luciferase levels quantified by IVIS bioluminescence imaging in the indicated treatment groups at day week 1 (W1) and week 3 (W3). Paired student’s t test. Ns, not significant. * p < 0.05. ** p < 0.01. *** p < 0.001. **** p < 0.0001.

